# Structural and functional analysis of YopR and identification of an additional key component of the SPβ phage lysis-lysogeny management system

**DOI:** 10.1101/2022.10.21.513154

**Authors:** Katharina Kohm, Ekaterina Jalomo-Khayrova, Syamantak Basu, Wieland Steinchen, Gert Bange, Robert Hertel, Fabian M. Commichau, Laura Czech

**Affiliations:** FG Synthetic Microbiology, Institute for Biotechnology, BTU Cottbus-Senftenberg, Senftenberg, Germany; FG Molecular Microbiology, Institute for Biology, University of Hohenheim, Stuttgart, Germany; Center for Synthetic Microbiology (SYNMIKRO) and Department of Chemistry, Phillips-University Marburg, Marburg, Germany; Max-Planck Institute for Terrestrial Microbiology, Marburg, Germany; Department of Genomic and Applied Microbiology, Institute of Microbiology and Genetics, Georg-August-University of Göttingen, Göttingen, Germany

**Keywords:** prophage, regulation, repressor, recombinase, suppressor mutant

## Abstract

Prophages need to tightly control their lifestyle to either be maintained within the host genome or enter the lytic cycle. The SPβ prophage present in the genome of *Bacillus subtilis* 168 was recently shown to possess an *arbitrium* system defining its replication stage. Using an historic *B. subtilis* strain harboring the heat-sensitive SPβ c2 mutant, we analyzed a key component of the lysis-lysogeny decision system called YopR, which is critical for maintenance of lysogeny. Here, we demonstrate that the heat-sensitive SPβ c2 phenotype is due to a single nucleotide exchange in the *yopR* gene, rendering the encoded YopR^G136E^ protein temperature sensitive. Structural characterization of YopR revealed that the protein is a DNA-binding protein with an overall fold like tyrosine recombinases. Biochemical and functional analyses indicate that YopR has lost the recombinase function and the G136E exchange impairs its higher order structure and DNA binding activity. We further show that the heat-inducible SPβ excision of the c2 mutant still depends on the serine recombinase SprA. Finally, an evolution experiment identified the YosL protein of unknown function as a novel component of the lysis-lysogeny management system, as the presence of *yosL* is crucial for the induction of the lytic cycle of SPβ.

## INTRODUCTION

Phages are bacterial viruses whose replication strictly depends on the host cell. While most host cells are killed after phage infection during the lytic cycle, some phages may enter the lysogenic or temperate life cycle [Salmond and Fineran, 2015; Dion et al., 2020]. In the latter case, the phage genome integrates into the host chromosome and the bacteria carrying the prophage become lysogens. Prophages multiplying together with the host chromosome may enter the lytic cycle through the action of mutagenic agents [Warner et al., 1977].

We are interested in the biology of the temperate phage SPβ that infects the endospore-forming Gram-positive model bacterium *Bacillus subtilis* and resides as a prophage in the genome of the *B. subtilis* laboratory strain 168 [Zeigler et al., 2008]. SPβ resembles the *Siphoviridae* morphotype and carries a 130 kb long genome [Kohm and Hertel, 2021]. It was discovered about 50 years ago in the *B. subtilis* strain CU1050 that was subjected to chemical mutagenesis and cured of the prophage [Georgopoulos, 1969; Warner et al., 1977; Johnson and Grossman, 2016]. Interestingly, cultures of *B. subtilis* that are lysogenic for SPβ release the *S*-linked glycopeptide sublancin, which belongs to the class of antimicrobial natural products named glycocins [Hemphill et al., 1980; Oman et al., 2011]. Sublancin inhibits the growth of SPβ-free *B. subtilis* strains by interfering with essential cellular processes such as DNA replication and transcription [Hemphill et al., 1980; Wu et al., 2019]. *B. subtilis* strains carrying the SPβ prophage in their genomes can therefore be identified based on their antimicrobial properties [Hemphill et al., 1980]. Like other temperate phages, SPβ may enter either the lytic or the lysogenic cycle. The lytic cycle of SPβ can be induced by treating *B. subtilis* with either *N*-methyl-*N’*-nitro-*N*-nitroso-guanidine or mitomycin C [Warner et al., 1977]. A study focusing on sporulation revealed that the SPβ genome is excised from the mother cell genome during the formation of the spore (Figure 1) [Abe et al., 2014]. The excision process depends on the serine recombinase SprA (<u>SP</u>β site-specific <u>r</u>ecombination factor <u>A</u>), whose activity is controlled by the accessory factor SprB [Abe et al., 2014, 2017, 2020]. Remarkably, the excision of the SPβ genome from the host cell genome causes the reconstitution of the *spsM* gene in the mother cell, which encodes an enzyme involved in sugar decoration of the spore and is thus crucial for spore maturation [Abe et al., 2014, 2017, 2020]. While the *sprA* gene is actively transcribed during vegetative growth, the expression of the *sprB* gene depends on the sporulation-specific sigma factors SigE and SigK [Abe et al., 2014]. The temporal synthesis of SprB prevents excision of the SPβ genome during vegetative growth of *B. subtilis*. Interestingly, while SPβ resides as a prophage in the spore genome, it is only excised in the mother cell to form the functional *spsM* gene. Moreover, this excision does not result in the formation of infectious phage particles [Abe et al., 2014].

**Figure 1.**
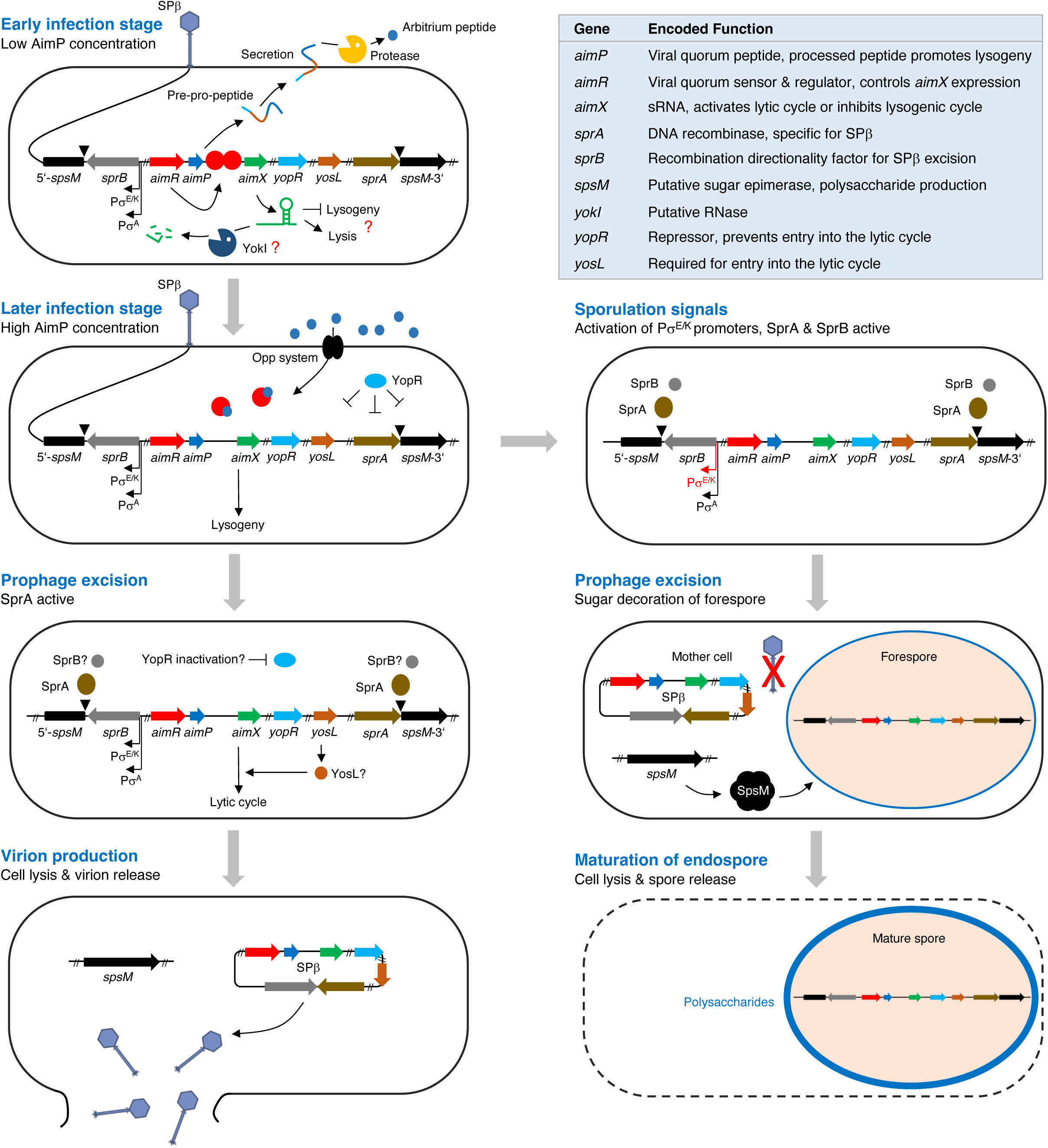
Major steps of the lytic and lysogenic cycle of SPβ. Gene and protein sizes are not drawn to scale. The target of the *aimX* sRNA is unknown and the potential RNase function of YokI remains to be elucidated. The signal that inhibits DNA-binding activity of YopR and the recombinase catalyzing the excision of the prophage are unknown.

Recently, it has been demonstrated that the phages of the SPβ group use a small-molecule communication system (termed the “arbitrium” system) to coordinate lysis-lysogeny decisions (Figure 1) [Erez et al., 2017; Trinh & Zeng, 2019]. The core of the arbitrium system consists of the *aimP*, *aimR* and *aimX* genes encoding the AimP peptide, the transcriptional activator AimR and the non-coding (nc) RNA AimX, respectively [Erez et al., 2017]. In the early stage of infection, the number of phages and the concentration of AimP is low. Apo-AimR binds as a homodimer *via* its helix-turn-helix domain to the operator in the *P_aimX_* promoter and activates transcription of the AimX ncRNA that promotes the lytic cycle (Figure 1) [Erez et al., 2017; Gallego Del Sol et al., 2019, 2022; Guan et al., 2019; Pei et al., 2021]. During the lytic cycle, accumulating AimP is secreted *via* the Sec pathway and processed by extracellular proteases. The resulting hexapeptide is transported into the cell by the Opp oligopeptide transport system [Lazzazera et al., 1997]. The formation of an AimP-hexapeptide/AimR complex inhibits the DNA-binding activity of AimR and promotes the switch to the lysogenic cycle [Erez et al., 2017]. While the molecular details of the AimP-hexapeptide/AimR complex have been elucidated [Wang et al., 2018; Dou et al., 2018; Zhen et al., 2019; Gallego Del Sol et al., 2019, 2022; Guan et al., 2019; Pei et al., 2021], the components that act downstream of the AimX ncRNA remain to be identified (Figure 1). However, the AimP-hexapeptide-dependent control of AimR is crucial to prevent the killing of the entire bacterial population by SPβ, thereby ensuring long-term prophage replication.

Two recent studies uncovered several novel players of the SPβ lysis-lysogeny decision system [Brady et al., 2021; Kohm et al., 2022]. For instance, a suppressor analysis with a SPβ *aimR* mutant, which generates more lysogenic cells, revealed that the spontaneous inactivation of the *yopN* gene, encoding a protein of unknown function, partially restores the control of lysis-lysogeny [Brady et al., 2021]. The same studies identified YopR as the master repressor of the lytic cycle of SPβ [Brady et al., 2021; Kohm et al., 2022]. Although it is unclear how AimR, YopN and YopR interact with each other, it has been proposed that YopN stimulates the repressor activity of YopR [Brady et al., 2021]. Furthermore, a recent comparative genome analysis identified the core genes of SPβ-like phages. These genes are *yopR*, *yopQ*, *yopP*, *aimR* and *yokI*, which are most likely transcriptionally active during the lysogenic cycle [Kohm et al., 2022]. The inactivation of the *yokI* gene indeed verified its role in the lysis-lysogeny decision system of SPβ because the *yokI* mutant released about four times more virions into the supernatant as compared to the parental strain [Kohm et al., 2022]. It has been hypothesized that YokI could act as an RNase that prevents the accumulation of the AimX ncRNA, thereby hindering the entry into the lytic cycle (Figure 1) [Kohm et al., 2022].

The above-mentioned prophage-free *B. subtilis* strain CU1050 also served as a host to isolate the “historic” SPβ mutant variants c1 and c2 that likely acquired mutations in genes of the lysis-lysogeny decision system [Zahler et al., 1977; Rosenthal et al., 1979]. The c1 mutant had an apparent plaque phenotype and did not lysogenize *B. subtilis*. By contrast, the c2 mutant proved to be temperature sensitive and entered the lytic cycle when the temperature was shifted to 50 °C for a few minutes. It has been suggested that c1 and c2 are alleles of the same gene encoding a repressor of SPβ like other temperate phages [Rosenthal et al., 1979]. However, the mutations in the SPβ c1 and c2 mutants and their impact on the lysogeny management system have not been identified and studied to date [McLaughlin et al., 1986]. Previous searches for regulator binding sites identified the <u>SPb</u>eta <u>r</u>epeated <u>e</u>lement (SPBRE) [Lazarevic et al., 1999]. Yet, the protein capable of binding to SPBRE has not been discovered.

Here, we demonstrate that the heat-sensitive SPβ c2 phenotype is due to a single nucleotide exchange in the *yopR* gene, rendering the encoded YopR^G136E^ protein temperature sensitive. Genetic complementation studies revealed that the wild type YopR protein is dominant over the YopR^G136E^ mutant. Thermal shift assays and biochemical analyses uncovered that the G136E exchange renders YopR less stable and reduces its affinity to bind DNA *in vitro*. Structural and functional characterization of YopR revealed that the protein is a DNA-binding protein that has an overall fold like tyrosine recombinases but lacks crucial amino acid residues to function as a recombinase enzyme. Moreover, cell lysis occurred after heat shock in strains lacking either serine recombinase SprA or its accessory factor SprB. Finally, a genetic suppressor analysis with a *B. subtilis* strain carrying the SPβ c2 *yopR* G407A allele identified the YosL protein as a novel player in the lysis-lysogeny management system. Further analysis showed that the presence of *yosL* seems to be crucial for the lytic cycle of SPβ. The current model of the SPβ lysis-lysogeny management system is discussed.

## MATERIALS AND METHODS

### Reagents

Primers used in this study were purchased from Sigma-Aldrich and are listed in Table S1. Chromosomal DNA was isolated from *B. subtilis* using the peqGOLD Bacterial DNA Kit (VWR). Plasmid DNA was isolated using the Nucleospin Extract Kit (Macherey-Nagel), Gene Jet Mini Prep Kit (Thermo Scientific), or Monarch® Plasmid Miniprep (NEB). DNA fragments that were generated by the polymerase chain reaction (PCR) were purified using the PCR Purification Kit (Qiagen), Monarch® PCR & DNA Cleanup Kit (NEB), or Gene Jet gel extraction kit (Thermo Scientific). Phusion DNA polymerase, restriction enzymes and T4 DNA ligase were purchased from Thermo Scientific, NEB, or Promega and used according to the manufacturer’s instructions [Sambrook et al., 1989]. Miscellaneous chemicals and media were purchased from Sigma-Aldrich, Carl Roth, and Becton-Dickinson. Plasmids were sequenced by the SeqLab Sequence Laboratories (Microsynth).

### Bacterial strains, media and growth conditions

Bacterial strains are listed in Table S2. *Escherichia coli* strains were grown in lysogeny broth (LB) at 37 °C with constant shaking at 200 rpm or on LB agar plates. Unless otherwise stated, *B. subtilis* was grown in LB or sporulation (SP) medium at 37 °C [Commichau et al., 2007]. For phage induction, fresh LB was inoculated to an OD_600_ of 0.1 and the culture was incubated for 2.5 h at 37 °C. A heat shock was then applied in a water bath for 10 min at 50 °C. To allow phage replication, the culture was further incubated for 2 h at 37 °C.

**Table 2.** Data collection and refinement statistics for YopR.

|  |  | YopR<br>(PDB code: 8a0a) |
| --- | --- | --- |
| **Data collection** |  |  |
| Space group |  | P2<sub>1</sub>2<sub>1</sub>2<sub>1</sub> |
| Cell dimensions |  |  |
|  |  | 49.55 |
| *a, b, c* (Å) |  | 98.87 |
|  |  | 162.24 |
|  |  | 90 |
| $\alpha, \beta, \gamma$ (°) | | 90 |
|  |  | 90 |
| Wavelength (Å) |  | 0.976253 |
| Resolution (Å) |  | 47.45 - 2.902 |
|  |  | (3.006 - 2.902) |
| *R*<sub>merge</sub> |  | 0.3707 (2.446) |
| *I*/$\sigma$(*I*) | | 8.89 (1.47) |
| Completeness (%) |  | 99.88 (99.27) |
| Redundancy |  | 12.4 (12.7) |
| *CC*<sub>1/2</sub> |  | 0.991 (0.581) |
| **Refinement** |  |  |
| Resolution (Å) |  | 47.45 - 2.902 |
| No. reflections |  | 226928 (22388) |
| *R*<sub>work</sub> / *R*<sub>free</sub> |  | 0.2149 / 0.2628 |
| No. atoms |  | 5182 |
| protein |  | 5182 |
| Ligands/ion |  | 0 |
| water |  | 0 |
| *B*-factor |  |  |
| protein |  | 67.98 |
| R.M.S. deviations |  |  |
| Bond lengths (Å) |  | 0.004 |
| Bond angles (°) |  | 0.75 |
| Ramachandran |  |  |
| favored (%) |  | 97.33 |
| allowed (%) |  | 2.67 |
| outliers (%) |  | 0.00 |

\*Statistics for the highest-resolution shell are shown in parentheses.
Data were collected on ID30B (ESRF).

### DNA manipulation, transformation, strain construction and phenotypic analysis

The *E. coli* strains XL1-Blue (Stratagene) and DH10B [Grant et al., 1990] were used for cloning and transformants were selected on LB agar plates supplemented with ampicillin (100 µg/ml) (Sigma-Aldrich or Carl Roth). *B. subtilis* was transformed with plasmid or chromosomal DNA according to the two-step protocol described previously [Kunst and Rapoport, 1995]. Transformants were selected on LB plates containing chloramphenicol (5 µg ml^-1^), kanamycin (25 µg ml^-1^), or erythromycin (5 µg ml^-1^). The plasmids constructed in this study are listed in Table S2. The plasmid pRH005 was constructed for the deletion of the *cat* gene in the strain TS01 using the CRISPR-Cas9 system [Altenbuchner et al., 2016]. The gRNA was introduced into the plasmid pJOE8999 using the primer pair RH080/RH081 resulting in plasmid pRH004. Next, DNA fragments flanking the *cat* gene were introduced into the plasmid pRH004 using the primer pairs RH082/RH083 and RH084/RH085. The deletion of the *cat* gene in the strain TS01 was performed as described previously [Schilling et al., 2018]. To enable the deletion of the genes of interest with the *ermC* deletion cassette [Koo et al., 2017], we had to remove the *ermD* resistance cassette in the *B. subtilis* strains KK001 and KK137, giving the strains KK002 (SPβ c2) and KK009 (SPβ), respectively. The plasmid pRH29 was generated for this purpose. The gRNA targeting the *ermD* gene was made by hybridizing the primers RH075 and RH076. The dsDNA fragment was ligated to pJOE8999 [Schilling et al. 2018; Otte et al. 2020] generating pRH001. The *ermD* deletion cassette was made with the primer pairs RH070/RH112 (flank A) and RH113/RH114 (flank B) and introduced into pRH001, resulting in pRH029. To express the *yopR*^G407A^ (YopR^G136E^), *yopR*^A911T^ (YopR^Y304F^) and *yopR*^TAT910-912GCG^ (YopR^Y304A^) mutant alleles from the *amyE* locus in *B. subtilis*, the plasmids pRH166, pRH180 and pRH181, respectively, were constructed. The *yopR*^G407A^, *yopR*^A911T^ and *yopR*^TAT910-912GCG^ alleles were amplified by PCR using the primer pair PP375/PP376, digested with *Eco*RI and *Bam*HI and ligated to pAC7 that was cut with the same enzymes. SPβ c2 DNA (*yopR*^G407A^) as well as the plasmids pEJK27 (*yopR*^A911T^) and the plasmid pEJK29 (*yopR*^TAT910-912GCG^) served as templates. The isogenic plasmid pRH167 carrying the wild-type *yopR* allele was constructed previously [Kohm et al., 2022]. The plasmid pRH182 carrying the *yopR*^AAG505-507GCG^ allele (YopR ^K169A^) was constructed as follows. Two halves of the plasmid pRH167 were amplified with the primer pairs PP396/HLHR4 and PP397/HLHR16 and purified. The fragments were combined and digested with the enzyme *Bsa*I. The remaining plasmid pRH167 was removed by digesting the sample with *Dpn*I. The purified fragments were then ligated and used to transform *E. coli* XL1-Blue. The integrity of the resulting plasmid pRH182 was verified by control digestion and Sanger sequencing. The expression of the *yopR* alleles from the plasmids pRH166, pRH167, pRH180, pRH181 and pRH182 is driven by the artificial *P_alf4_* promoter and the ribosome-binding site of the *B. subtilis gapA* that were attached by PCR [Gundlach et al., 2017]. To express the wild type *yopR* and the *yopR^G407A^* alleles from the *ganA* locus in *B. subtilis*, we constructed the plasmid pRH174 and pKK005, respectively. The *yopR* alleles were amplified by PCR using the primer pair PP375/PP376 and introduced into the plasmid pGP888 via the enzymes *Eco*RI and *Bam*HI. The artificial *P_alf4_* promoter for constitutive expression was attached by PCR. To study the role of YopR on the activity of the *P_aimR_* promoter, we constructed the plasmid pRH173 (*P_aimR_ cat*). For this purpose, we amplified the intergenic region upstream of the *aimR* gene with the primer pair and PP390/PP391 and introduced the fragment into the plasmid pAC5 using the restriction sites of *Eco*RI and *Bam*HI. Plasmid pRH168 was constructed for the xylose-dependent *yosL* gene from the *ganA* locus. For this purpose, the *yosL* gene was amplified by PCR with the primer pair PP366/PP367, digested with *Eco*RI and *Bam*HI and ligated to pGP888 that was cut with the same enzymes. The SPβ and SPβ c2 phages were isolated from overnight cultures of the *B. subtilis* strains 168 and CU1147, respectively, by centrifuging 2 ml of the cultures for 1 min at 10,000 x g. Bacterial cells were removed by filtration and the sterile supernatants were used to infect the *B. subtilis* strain TS01 resulting in the lysogens KK001 (SPβ) and KK137 (SPβ c2). The isolation of lysogens from plaques and the sublancin assay to verify the presence of the prophages were described previously [Otte et al., 2020; Kohm et al., 2022]. For the purification of the YopR, YopR^G136E^ and YosL proteins, the underlying gene fragments were amplified by PCR using the primer pairs LC195/LC196 for both versions of YopR and LC192/193 for YosL and introduced into a pET24d (Novagen) derivative modified for modular cloning via BsaI restriction sites. This resulted in the plasmids pEJK11 (YopR), pEJK13 (YopR^G136E^) and pEJK9 (YosL) (Table S2). All expression constructs contained a C-terminal hexa-histidine (His_6_) tag fused to the target protein. Proteins derived from *B. subtilis* were produced in *E. coli* BL21 (DE3) (NEB). Deletion of the *sprA* and *sprB* in the strain KK002 was achieved by transformation using PCR products that were generated with the primer pairs PPKK001/PPKK002 and PPKK003/PPKK004, respectively, and chromosomal DNAs of the strains BKK21660 and BKK19820 as the template.

### Site-directed mutagenesis

For the overproduction of YopR variants with amino acid substitutions, we constructed the plasmids pEJK25 (YopR^K169A^), pEJK27 (YopR^Y304F^), and pEJK29 (YopR^Y304A^) using the Q5 Site-Directed Mutagenesis Kit (New England Biolabs; Ipswich, USA) according to the manufacturer’s protocol. The primers for mutagenesis were designed with the NEBaseChanger online tool and are listed in Table S1.

### Genome sequencing

Genomic DNA was prepared from 500 µl overnight cultures using the peqGold bacterial DNA mini kit following the instruction of the manufacturer with the modification of physically opening cells with the TissueLyser II (Qiagen). Purified genomic DNA was paired end sequenced (2 x 150 bp) (GENEWIZ). Sequence libraries were prepared with the NEBNext Ultra II FS DNA Library Prep 136 kit (New England Biolabs GmbH, Frankfurt, Germany) and sequenced with the NovaSeq6000 (Illumina, San Diego, CA, USA). The sequencing reads were mapped onto the genome of the *B. subtilis* strain SP1 and mutations were identified using the applications bowtie2 and breseq [Langmead and Salzberg, 2012; Deatherage and Barrick, 2014]. SP1 is prototrophic for tryptophan and a derivative of the laboratory strain 168 [Richts et al., 2020]. All identified mutations were verified by PCR and Sanger sequencing.

### Electrophoretic mobility shift assays

EMSAs were carried out to analyze the DNA-binding activity of wild type YopR and its variants. Regions containing the P*_aimR_*, P*_yosX_* and *attR* sites were amplified from *B. subtilis* WT168 chromosomal DNA using the primer pairs LC226/LC252, LC230/LC254 and LC255/LC256, respectively (Table S1). In a binding reaction, 1 pmol of the DNA fragment was mixed with the indicated protein concentrations in EMSA buffer containing 20 mM HEPES-Na (pH 7.5), 200 mM NaCl, 20 mM KCl, 50 µg ml^−1^ herring sperm DNA in a final volume of 20 µl. After incubation of the reaction mixture at RT for 15 min, samples were loaded onto a 2% (w/v) agarose gel (in 1 × TBE containing 90 mM Tris, 90 mM boric acid, and 2 mM EDTA). Samples were separated at 100 V for 90 min, stained in 1 x TBE buffer containing ethidium bromide and subsequently imaged using a UV detector system.

### *In vitro* recombination assays

The *in vitro* recombination assays were performed as described previously [Abe et al., 2017] with minor modifications. Fluorescently labeled DNA substrates were amplified from *B. subtilis* wild type strain 168 chromosomal DNA using the primer pairs LC255*(Cy3 modification at 5’)/LC256 (610 bp) for *attR* site and EJK54/EJK55*(Cy5 modification at 5’) (242 bp) for *attL* site (Table S1). In the assayed reaction, 20 nM of the DNA substrates were mixed with 20 µM YopR or the indicated YopR variant in a buffer containing 20 mM HEPES-Na (pH 7.5), 200 mM NaCl and 20 mM KCl, in a final volume of 10 µl. The mixture was incubated at 37 °C for 60 minutes. The recombination reaction was stopped by the addition of 0.1% (w/v) SDS and by heat treatment at 85 °C for 3 min. Samples were loaded onto a native 6% (w/v) polyacrylamide gel (in 0.5 × TBE containing 45 mM Tris, 45 mM boric acid and 1 mM EDTA). Samples were separated at 100 V for 90 min and subsequently imaged with the 600 and 700 nm channels of an Odyssey FC Imager (LI-COR Biosciences, Lincoln, US).

### β-galactosidase activity assay

Quantitative studies of *lacZ* expression in *B. subtilis* were performed as described previously [Kunst and Rapoport, 1995]. Cells were grown at 37 °C and 220 rpm in 30 ml of LB medium. The medium was supplemented in 300 ml shake flasks and the cells were grown without additional aeration. Cells were harvested before heat shock and at, 30 min, 60 min, 90 min, 120 min and 150 min incubation at 50 °C. Specific β-galactosidase activities were determined with cell extracts obtained by lysozyme treatment. One unit of β-galactosidase is defined as the amount of enzyme which produces 1 nmol of *o*-nitrophenol per min at 28 °C. The BioRad dye-binding assay was used to determine the protein concentrations.

### Overexpression and protein purification

For recombinant overexpression of YopR-His_6_ (pEJK11), the YopR-His_6_ variants G136E (pEJK13), K169A (pEJK25), Y304F (pEJK27) and Y304A (pEJK29) and YosL-His_6_ (pEJK9), 2 l of LB medium containing kanamycin (50 µg ml^-1^) and 1% (w/v) lactose for autoinduction of the *P_lac_* promoter driving the expression of the T7 polymerase required for recombinant gene expression were incubated in an aerial shaker for 18 h at 30 °C. After harvesting, cells were lysed by a microfluidizer (M110-L, Microfluidics). The lysis buffer contained 20 mM HEPES-Na (pH 8.0), 250 mM NaCl, 20 mM KCl and 40 mM imidazole. Cell debris was then removed by high-speed centrifugation for 20 min at 48,000 × g. All proteins were initially purified by nickel ion affinity chromatography and eluted with lysis buffer containing 250 mM imidazole. Further purification using HiTrap Heparin HP columns (Cytiva, Marlborough, USA) was performed according to the manufactures protocol. The eluted proteins were concentrated by centrifugation (3 kDa or 10 kDa MWCO) and further polished by size-exclusion chromatography on a S200 XK16 column (for YopR and variants) or S75 XK16 column (for YosL) (Cytiva, Marlborough, USA) with size-exclusion chromatography (SEC) buffer consisting of 20 mM HEPES-Na (pH 7.5), 200 mM NaCl and 20 mM KCl. Purified proteins were analyzed for the presence of bound nucleotides by agarose gel electrophoresis and analytical size-exclusion chromatography.

### Analytical size-exclusion chromatography

For the analytical SEC, purified YopR and YopR variants were diluted in a buffer containing 20 mM HEPES-Na (pH 6.37), 200 mM NaCl and 20mM KCl to a final concentration of 100 µM. 100 µl were then injected at 4 °C on to a pre-equilibrated S200 300/10 GL analytical size-exclusion column (Cytiva, Marlborough, USA) on an ÄKTA system (UNICORN 7.6; Cytiva). For size calibration, a mixture of standard proteins [thyroglobulin (660 kDa), ferritin (474 kDa), aldolase (160 kDa), conalbumin (76 kDa), ovalbumin (43 kDa) and ribonuclease A (13.7 kDa)] was used according to the manufacturer’s protocol (Cytiva, Marlborough, USA). Data were plotted using GraphPad Prism (GraphPad Prism Corp. San Diego, USA).

### Thermal shift assays using nanoDSF

To assess the protein folding and thermal stability of YopR and its variants, thermal shift assays were performed with nanoDSF integrated in the Prometheus NT.48 (NanoTemper Technologies GmbH, Germany). Protein solutions of 75-100 µM of YopR WT, YopR^G136E^, YopR^K169A^, YopR^Y304A^ and YopR^Y304F^ were soaked into a Prometheus NT.48 Series nanoDSF Grade Standard Capillaries and analyzed using PR. ThermControl software. Data were evaluated and plotted using GraphPad Prism (GraphPad Prism Corp., San Diego, USA). The normalized data were fitted to Boltzmann sigmoidal equation. The T_m_ values correspond to the inflection point of the sigmoidal curve [Boltzmann, 1970; Niesen et al., 2007].

### Isothermal titration calorimetry

Ligands and proteins (purified wild type protein YopR and YopR^G136E^ and YopR^Y304F^ variants) were diluted with a buffer containing 20 mM HEPES-Na (pH 7.5), 200 mM NaCl and 20 mM KCl. The DNA ligand (*aimR* promotor) was obtained by the hybridization of LC292/LC293 primers at 95 °C for 2 minutes. The sample cell was filled with DNA (*aimR* promotor) at a nominal concentration of 20 µM. The YopR proteins were placed in the syringe and their concentrations were predetermined by absorbance at 280 nm to saturate the DNA sample during the titrations. All the measurements were performed at 25 °C with the instrument MicroCal PEAQ-ITC (©Malvern Panalytical) with a method consisting of 13 injections (first 0.4 µl, and the rest 3 µl each) and 150 s of spacing. The raw data were processed with the MicroCal PEAQ-ITC Analysis Software (Malvern Panalytical) using the “one set of sites” models.

### Crystallization and structure determination

Crystallization was performed by the sitting drop vapor diffusion method at 20 °C in 250 nl drops consisting of equal parts of protein and precipitation solutions. Protein solutions of 480 µM YopR-His_6_ were used for crystallization. A total of 384 different conditions were included in the screen (JCSG Core Suite I-IV, Qiagen) and crystals obtained in 0.2 M disodium tartrate supplemented with 20% (w/v) PEG3350. Prior to data collection, crystals were flash-frozen in liquid nitrogen using a cryo-solution that consisted of mother liquor supplemented with 20% (v/v) glycerol. Data were collected under cryogenic conditions at the European Synchrotron Radiation Facility (Grenoble, France) [Theveneau et al., 2013]. MxCube3 was used for data collection (https://github.com/mxcube). Data were processed with XDS (version January 31, 2020) and scaled with XSCALE [Kabsch, 2010]. The structure was determined by molecular replacement with PHASER [McCoy, 2006], manually built in COOT (Coot Version 0.9.4.1) [Emsley and Cowtan, 2004], and refined with PHENIX [Liebschner et al., 2019] (Phenix Version 1.17.1-3660 and 1.19). The search model for the YopR structure was the N-terminal domain (residues 1-105) of the Alphafold2 model [Jumper et al., 2021]. Figures were prepared with Pymol (www.pymol.org). Crystallization data collection and refinement statistics are given in Table 2. Structure coordinates and structure factors of YopR have been deposited under the PDB-ID: 8a0a (https://doi.org/10.2210/pdb8A0A/pdb).

### Hydrogen/deuterium exchange mass spectrometry

Preparation of samples for HDX-MS was aided by a two-arm robotic autosampler (LEAP Technologies) as described previously with minor modifications [Osorio-Valeriano et al., 2019]. YopR protein variants and DNA (where indicated) were employed at concentrations of 25 µM each. Hydrogen/deuterium exchange (HDX) was initiated by 10-fold dilution of YopR, YopR^G136E^ or YopR^Y304F^ in buffer (20 mM HEPES-Na pH 7.5, 20 mM KCl, 200 mM NaCl) prepared in D_2_O. After incubation at 25 °C for 10, 30, 100, 1,000 or 10,000 s, the HDX was stopped by mixing the reaction with an equal volume of quench buffer (400 mM KH_2_PO_4_/H_3_PO_4_, 2 M guanidine-HCl; pH 2.2) temperated at 1 °C, and 100 μl of the resulting mixture injected (loop volume 50 µl) into an ACQUITY UPLC M-Class System with HDX Technology [Wales et al., 2008]. Non-deuterated samples were generated similarly by 10-fold dilution in buffer prepared with H_2_O. The injected HDX samples were washed out of the injection loop with H_2_O + 0.1% (v/v) formic acid at 100 µl min^-1^ flow rate and guided over a column (2 mm x 2 cm) filled with immobilized protease facilitating proteolytic digestion at 12 °C. The resulting peptides were collected on a trap column (2 mm x 2 cm), that was filled with POROS 20 R2 material (Thermo Scientific) kept at 0.5 °C. After three minutes of digestion and trapping, the trap column was placed in line with an ACQUITY UPLC BEH C18 1.7 μm 1.0 x 100 mm column (Waters) and peptides eluted through a gradient of H_2_O + 0.1% (v/v) formic acid (eluent A) and acetonitrile + 0.1% (v/v) formic acid (eluent B) at 60 μl min^-1^ flow rate as follows: 0-7 min/95-65% A, 7-8 min/65-15% A, 8-10 min/15% A. Eluting peptides were guided to a Synapt G2-Si mass spectrometer (Waters) and ionized with by electrospray ionization (250 °C capillary temperature, 3.0 kV spray voltage). Mass spectra were acquired from 50 to 2,000 *m/z* in enhanced high-definition MS (HDMS^E^) [Geromanos et al., 2009; Li et al., 2009] or high-definition MS (HDMS) mode for non-deuterated and deuterated samples, respectively. Continuous lock mass correction was implemented with [Glu1]-Fibrinopeptide B standard (Waters). During separation of the peptides on the ACQUITY UPLC BEH C18 column, the pepsin column was washed three times by injecting 80 μl of 0.5 M guanidine hydrochloride in 4% (v/v) acetonitrile. Blank runs (injection of H_2_O instead of protein) were performed between each sample. The experiment was conducted twice in triplicates (individual HDX reactions) whereby either immobilized porcine pepsin or a mixture of immobilized protease type XVIII from *Rhizopus* sp. and protease type XIII from *Aspergillus saitoi* were employed for proteolytic digestion. Peptides were identified and evaluated for their deuterium incorporation with the softwares ProteinLynx Global SERVER 3.0.1 (PLGS) and DynamX 3.0 (both Waters) as described [Osorio-Valeriano et al., 2019]. Peptides were identified with PLGS from the non-deuterated samples acquired with HDMS^E^ employing low energy, elevated energy and intensity thresholds of 300, 100 and 1,000 counts, respectively and matched using a database containing the amino acid sequence of YopR, porcine pepsin and their reversed sequences with search parameters as follows: Peptide tolerance = automatic; fragment tolerance = automatic; min fragment ion matches per peptide = 1; min fragment ion matches per protein = 7; min peptide matches per protein = 3; maximum hits to return = 20; maximum protein mass = 250,000; primary digest reagent = non-specific; missed cleavages = 0; false discovery rate = 100. For quantification of deuterium incorporation with DynamX, the data obtained from pepsin or fungal protease digestion were combined, and peptides had to fulfil the following criteria: Identification in at least 2 of 3 non-deuterated samples for either protease digestion protocol; the minimum intensity of 10,000 counts; maximum length of 30 amino acids; minimum number of products of two; maximum mass error of 25 ppm; retention time tolerance of 0.5 minutes. All spectra were manually inspected and omitted, if necessary, e.g. in case of low signal-to-noise ratio or the presence of overlapping peptides disallowing the correct assignment of the isotopic clusters. Residue-specific deuterium uptake from peptides identified in the HDX-MS experiments was calculated with the software DynamX 3.0 (Waters). In the case that any residue is covered by a single peptide, the residue-specific deuterium uptake is equal to that of the whole peptide. In the case of overlapping peptides for any given residue, the residue-specific deuterium uptake is determined by the shortest peptide covering that residue. Where multiple peptides are of the shortest length, the peptide with the residue closest to the peptide C-terminus is utilized. Raw data of deuterium uptake by the identified peptides and residue specific HDX are provided in Supplemental Dataset.

### Western blot analysis

Heterologously produced and purified YopR-His_6_ and YosL-His_6_ proteins were used to generate rabbit polyclonal antibodies (Dr. Benli, Göttingen). For Western blot analyses, the cells were grown as described for the β-galactosidase activity assay. Proteins were separated by 15% SDS-PAA gels. After electrophoresis, the proteins were transferred to a polyvinylidene difluoride membrane (PVDF, BioRad) by electroblotting. YopR and YosL were detected with polyclonal antibodies [Commichau et al., 2007]. Antibodies were visualized by using anti-rabbit immunoglobulin G-alkaline phosphatase secondary antibodies (Promega) and the CDP-star detection system (Roche Diagnostics) as described previously [Commichau et al., 2007].

## RESULTS

### Verification of the SPβ c2 phenotype and generation of lysogens in a prophage-free strain

To verify the heat-sensitive phenotype of the historical *B. subtilis* strain CU1147 carrying the SPβ c2 prophage, we performed a heat shock experiment [Zahler et al., 1977; Rosenthal et al., 1979]. For this purpose, cultures containing exponentially growing cells of CU1147 were shifted from 37 °C to 50 °C for 5 minutes as described previously [Rosenthal et al., 1979]. As shown in Figure 2A, only the cells of CU1147 started to lyse 1 hour after heat treatment, confirming the presence of the SPβ c2 in the genome of CU1147 and the heat-sensitive phenotype of the prophage. To explore the underlying genotype of the SPβ c2 phage mutant and to prevent a potential crosstalk between SPβ c2 and other prophages and prophage-like elements, we established the *B. subtilis* strain TS01 as the new host. The TS01 strain is a descendant of 168, lacking the prophages SPβ and PBSX, the prophage-like elements prophage 1, prophage 3, and *skin*, as well as the *pks* polyketide synthesis operon [Westers et al., 2003; Reuss et al., 2016; Schilling et al., 2018]. New lysogens carrying SPβ and SPβ c2 designated as KK009 and KK002, respectively, were isolated from plaques of TS01-infected cells (Figure 2B, upper panel). Further experiments revealed that a cultivation temperature of 37 °C and a 10 min-long heat shock are best suited to induce the lytic cycle of SPβ c2. The sublancin assay indicated the presence of the prophages and the subsequent heat shock experiment confirmed that the heat-induced cell lysis was caused by SPβ c2 (Figure 2B, lower panel; Figure 2C).

**Figure 2.**
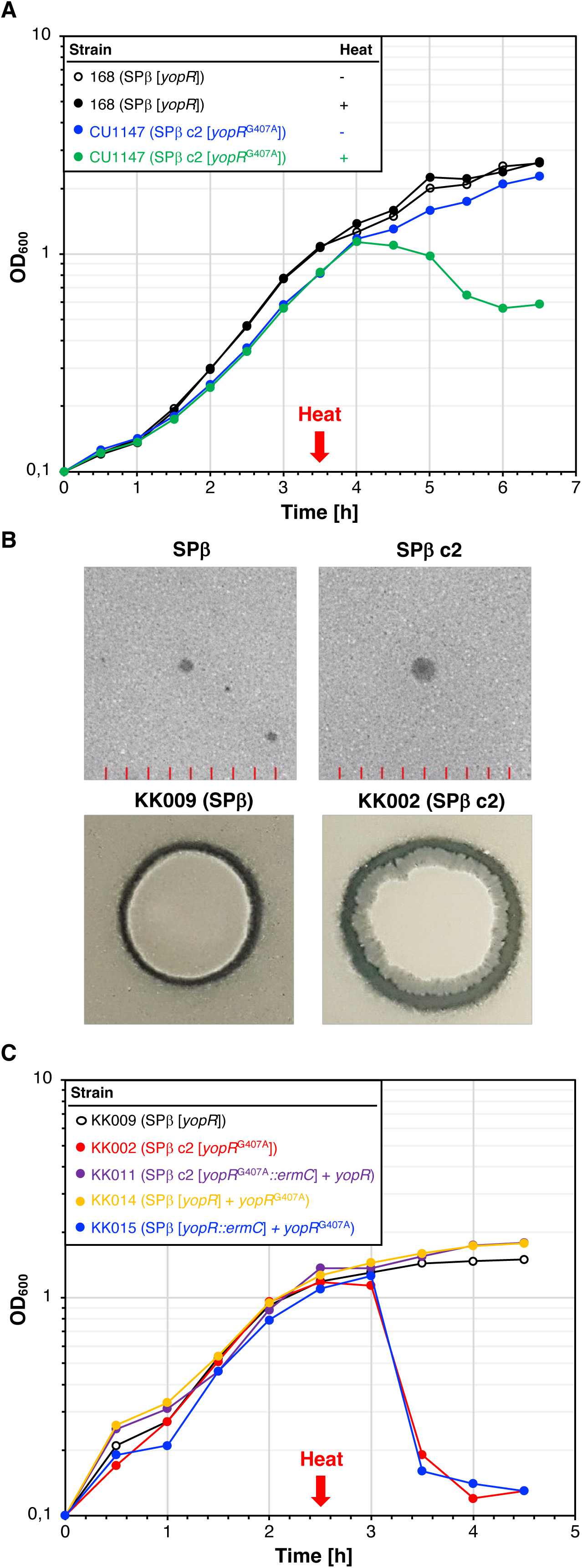
The temperature sensitive SPβ c2 prophage produces sublancin and senses heat shocks due to a mutation in the *yopR* gene. **A.** Heat shock experiment to assess the temperature sensitivity of the SPβ c2 prophage in the historical *B. subtilis* strain CU1147. Cultures of the strains 168 (SPβ [*yopR*], control) and CU1147 (SPβ c2 [*yopR*^G407A^]) were grown for 2.5 h at 30 °C, heat shocked at 50 °C for 10 min and further cultivated at 30 °C. The gene marked with square brackets is present in the phage genome. The optical density at a wavelength of 600 nm (OD_600_) was monitored over time. Further induction experiments revealed 37 °C as growing temperature and 10 min heat shock as best to induce SPβ c2. **B.** Plaque assay (upper panel) with SPβ and SPβ c2 lysates and sublancin assay with the strains KK009 and KK002 carrying SPβ and SPβ c2 prophages, respectively (see Materials and Methods). **C.** Heat shock experiment showing that the *yopR*^G407A^ allele causes the temperature-sensitive phenotype of the SPβ c2 prophage. The indicated strains were heat shocked as described in the Materials and Methods section. The gene marked with square brackets is present in the phage genome. The OD_600_ was monitored over time. The strains KK011, KK014 and KK015 express the *yopR/yopR*^G407A^ alleles from the *amyE* locus.

### The YopR^G136E^ variant confers heat induction of the SPβ c2 prophage

Previously, it has been suggested that the c2 allele encodes a repressor of SPβ [Rosenthal et al., 1979]. However, the mutation causing the c2 phenotype remained unknown. To identify the mutation(s) conferring heat inducibility of SPβ c2, we re-sequenced the SPβ c2 genome. Sequence variations were identified through a direct alignment with the SPβ genome of the *B. subtilis* laboratory strain SP1 [Richts et al., 2020]. We identified four sequence variations in the *yorN*-*yorM* intergenic region and in the *yoqA*, *yopR* and *yokI* genes. The mutation in the *yorN*-*yorM* intergenic region could also affect either the transcription of the *yorN* gene of unknown function or the translation of the *yorN* mRNA. The sequence deviations in the remaining three genes cause amino acid exchanges in the encoded proteins (Table 1). While YoqA shares no similarity to any known protein domains, YokI and YopR share sequence similarities with ribonucleases and DNA breaking-rejoining proteins, respectively. Since it was recently shown that YopR is the master repressor of the lytic cycle of SPβ [Brady et al., 2021; Kohm et al., 2022], we speculated that the base exchange G407A in the coding region of *yopR* is responsible for the heat-sensitivity of SPβ c2. To test this idea, we fused the constitutively active synthetic *P_alf4_* promoter [Gundlach et al., 2017] and the ribosome binding site of the *B. subtilis gapA* gene to the *yopR* wild type and the *yopR*^G407A^ alleles and integrated the constructs into the *amyE* locus of the strains KK002 (SPβ c2 [*yopR*^G407A^]) and KK009 (SPβ [*yopR*]), respectively. The resulting strains were designated as KK011 (SPβ c2 [*yopR*^G407A^] *P_alf4_-yopR*) and KK014 (SPβ [*yopR*] *P_alf4_-yopR*^G407A^). Next, we constructed the strain KK015 (SPβ [*yopR::ermC*] *P_alf4_-yopR*^G407A^) lacking the native SPβ *yopR* allele. The following heat shock experiment confirmed that the heat-induced cell lysis of *B. subtilis* was due to the mutation in *yopR* because all strains carrying SPβ and the *yopR*^G407A^ allele (KK002 and KK015) lysed after the heat shock (Figure 2C). Furthermore, no cell lysis was observed with the strains KK011 (SPβ c2 [*yopR*^G407A^*::ermC*] *P_alf4_-yopR*) and KK014 (SPβ [*yopR*] *P_alf4_-yopR*^G407A^) that carried both *yopR* alleles (Figure 2C). Thus, the wild type YopR protein is dominant over the YopR^G136E^ variant. To conclude, the G407A mutation in *yopR* causes the heat sensitivity of YopR^G136E^.

**Table 1.** Mutations identified in the SPβ c2 phage.

| Region | Encoded function | Type of mutation | Coordinates<sup>a</sup> |
| --- | --- | --- | --- |
| *yokI* | Putative ribonuclease | G145A (G49S) | 2.277.281 |
| *yopR* | Repressor | G407A (G136E) | 2.204.767 |
| *yoqA* | Unknown | T68C (L23P) | 2.201.412 |
| *yorN-yorM* | Unknown | ΔAG, 25-24 bp upstream of *yorN* | 2.174.763 - 2.174.762 |

<sup>a</sup>Coordinates refer to the position in the genome sequence CP058242 of the *B. subtilis* SP1 strain [Richs et al., 2020].

### Characterization of YopR and YopR^G136E^

To assess whether the heat shock affects the cellular concentration of the YopR^G136E^ variant, we performed a Western blot experiment with crude extracts from the strains TS01 (no phage) and KK002 (SPβ c2) that were grown over night at 37 °C. We also analyzed samples from a culture of the strain KK002 that was grown for 2.5 h at 37 °C and for 10 min at 50 °C. As expected, the strain TS01 did not synthesize YopR (Figure 3A). By contrast, the strain KK002 produced a protein corresponding to the molecular weight of YopR. Moreover, the bacteria produced comparable amounts of YopR^G136E^ variant before and after the heat shock. Thus, the G136E exchange in YopR probably does not affect the *in vivo* stability of the protein.

**Figure 3.**
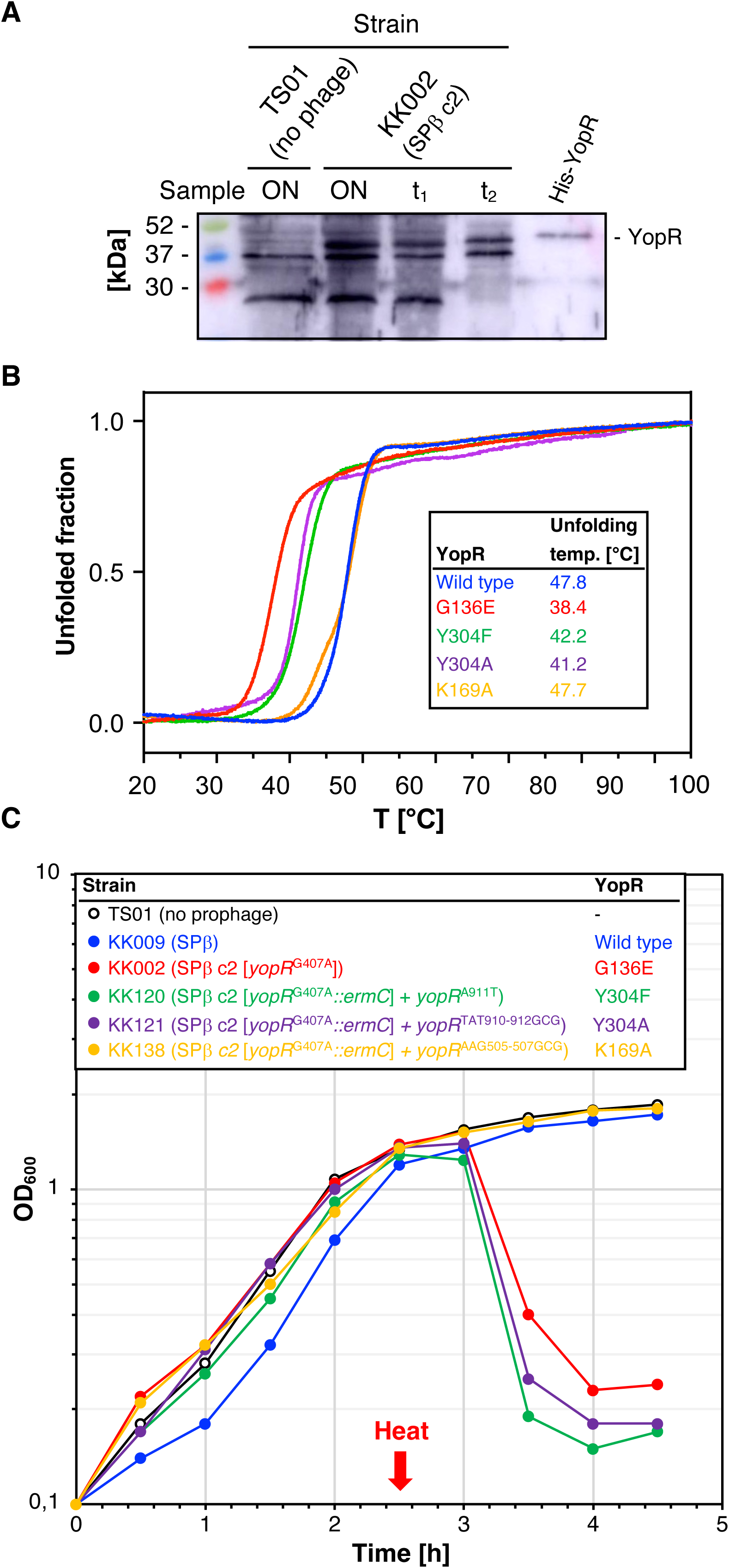
Cellular amounts of YopR ^G136E^ and characterization of YopR variants. **A.** Cellular amounts of YopR^G136E^ were determined by Western blotting. Crude extracts of the indicated strains were isolated from overnight (ON) or exponentially growing LB cultures at timepoints t_1_ and t_2_, immediately and 30 min after the heat shock (10 min incubation at 50 °C), respectively. As a control served purified YopR-His_6_ (100 ng). After electrophoresis in a 12.5% SDS-PAGE and transfer onto a PVDF membrane, YopR was detected by rabbit polyclonal antibodies raised against YopR-His_6_. 20 mg of crude extracts were applied. **B.** Analysis of purified YopR wild type (blue), YopR^G136E^ (red), YopR^Y304F^ (green), YopR^Y304A^ (purple) and YopR^K169A^ (orange) stability by nanoDSF. Thermal unfolding curves of YopR and YopR variants were assessed in SEC buffer. Changes in the F350/F330 fluorescence ratio are shown and T_m_ values are indicated in the table. **C.** Heat shock experiment to assess the temperature sensitivities of the YopR wild type (blue), YopR^G136E^ (red), YopR^Y304F^ (green), YopR^Y304A^ (purple), and YopR^K169A^ (orange). The indicated strains were heat shocked as described in the Materials and Methods section. The OD_600_ was monitored over time.

Next, we assessed the effect of the temperature on the folding of the YopR wild-type protein and the YopR^G136E^ variant *in vitro*. For this purpose, the affinity-purified proteins were assessed for their unfolding behavior in response to increasing temperature by nanoDSF. While the YopR wild-type protein exhibited an unfolding temperature of 47.8 °C, the heat-sensitive YopR^G136E^ variant unfolded approximately 10 °C earlier, at a temperature of 38.4 °C (Figure 3B). To assess the correct folding of YopR^G136E^, the variant was also analyzed by analytical size-exclusion chromatography (SEC). The wild-type protein as well as the YopR^G136E^ variant showed a single symmetrical peak corresponding to the size of a YopR monomer (calculated size: 36 kDa, experimentally determined size: 43.06 kDa for YopR-WT and 53.8 kDa for YopR^G136E^ (Figure S1A, C). Additionally, examination of the proteins through SDS-PAGE revealed signs of degradation for YopR^G136E^ (Figure S1G). Hence, mutation of G136E in YopR renders the protein less stable and temperature sensitive *in vitro*.

### YopR is a DNA binding protein

To further analyze the role of the G136E amino acid exchange in YopR, we aimed to study the protein biochemically. Hence, the YopR wild type protein and the YopR^G136E^ variant carrying a C-terminal His_6_-tag were produced in *E. coli* and purified by Ni-NTA affinity purification. During initial purification of YopR, co-purification of substantial amounts of DNA was observed, indicating that YopR is a DNA-binding protein (Figure S2). To obtain a DNA-free YopR protein sample an additional purification step using a Heparin column and a salt-gradient elution was performed after Ni-NTA purification (Figure S1A, B). Absence of DNA was assessed by analyzing the molecular weight of purified YopR by analytical size-exclusion chromatography (Figure S1A-C and G). We further wanted to test if YopR specifically binds to sequences found in the genome/intergenic regions of SPβ. Hence, electrophoretic mobility shift assays (EMSAs) using purified DNA-free YopR and YopR^G136E^ were performed with the fragments upstream of *aimR* and *yosX* harboring the previously described SPBRE (SPβ repeated element) [Lazarevic et al., 1999]. While the YopR wild type protein only required the presence of 5-fold protein excess to result in a shift of the DNA fragments, the YopR^G136E^ variant showed a decrease in binding affinity (approx. 10 to 20-fold excess of YopR^G136E^ required for a similar DNA shift) (Figure 4A-D). To further assess the reduced binding affinity, we performed binding studies using isothermal titration calorimetry (ITC). We employed the region upstream of the *aimR* promoter and added increasing amounts of the YopR wild type protein or the YopR^G136E^ variant. While the wild type protein had a DNA binding affinity of 0.238 ± 0.08 µM (Figure S3A, C), the binding affinity of the YopR^G136E^ variant was increased by approximately 7-fold to 1.69 ± 0.69 µM (Figure S3B, D). To conclude, the EMSAs and ITC analysis indicate that YopR is a DNA-binding protein and that the G136E exchange reduces its affinity for DNA. Since it was unclear if the tested DNA fragments containing the previously described SPBRE (<u>SP</u>β <u>r</u>epeated <u>e</u>lement) are the correct DNA binding targets of YopR, we also analyzed DNA binding of the *attR* region at the corner of the SPβ genome and an unrelated gene from *B. subtilis* (*yaaA*) (Figure 4E, F and Figure S4). EMSAs of both regions indicate a DNA binding behavior of the wild-type YopR protein like that observed for the SPBRE-containing regions upstream of the SPβ *aimR* and *yosX* genes (Figure 4E and Figure S4A). In contrast, the EMSA assessing the binding of YopR^G136E^ to the unrelated gene from *B. subtilis* (*yaaA*) indicates even less binding affinity as compared to the other tested regions. While for YopR^G136E^ a shift of the DNA-protein complex using the *attR* region, and the regions upstream of the SPβ *aimR* and *yosX* genes was observed upon addition of approximately 20-fold protein excess, a shift of the *yaaA* genetic region was only observed in a DNA:protein-ratio of 1:100 (Figure S4B). Therefore, the factors that establish the specificity of YopR DNA binding are still unclear.

**Figure 4.**
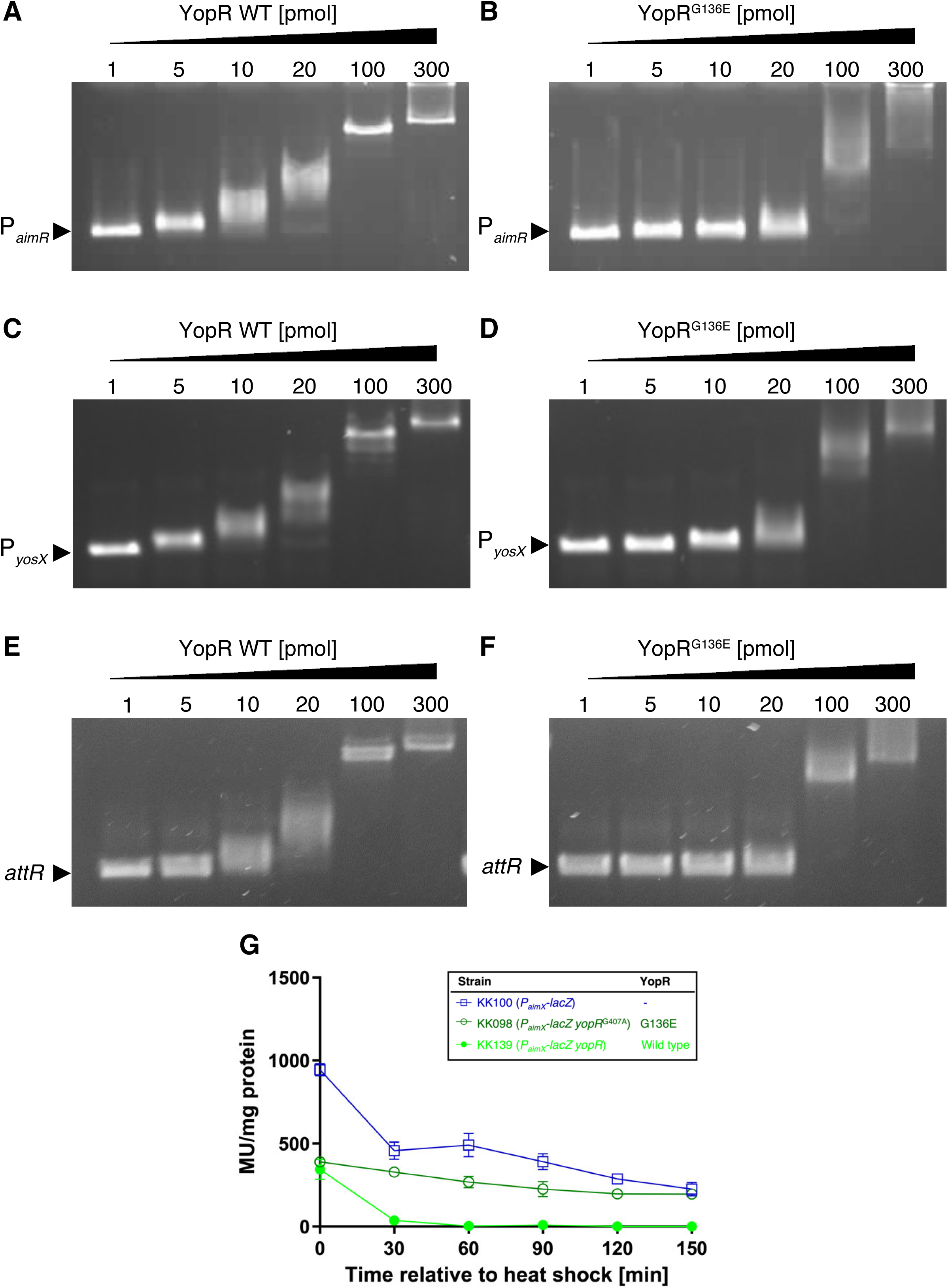
In vitro and in vivo analysis of YopR DNA binding. **A-F.** Electrophoretic mobility shift assays (EMSAs) showing the DNA-binding activities of the YopR wild type protein and the YopR^G136E^ variant**. A, C, F.** Complex formation between the YopR wild-type protein and the *P_aimR_, P_aimX_ and attR* DNA fragments. **B, D, F.** Complex formation between the YopR^G136E^ variant and the *P_aimR_, P_aimX_ and attR* DNA fragments. See Materials and Methods for the experimental procedure. **G.** β-Galactosidase activity assays to assess the DNA-binding activity of wild type YopR and YopR^G136E^. The indicated strains were grown in LB medium for 2.5 h at 37 °C, transferred to 50 °C and further cultivated for 2.5 h. The β-galactosidase activity is given as units per milligram protein. Experiments were carried out at least two times. Boxes indicate mean values.

### *In vivo* DNA binding activity of YopR

To assess the effect of the environmental temperature on the *in vivo* DNA-binding activity of YopR and YopR^G136E^, we monitored the activity of a translational *P_aimR_-lacZ* fusion. For this purpose, the *yopJ-aimR* (219 bp) intergenic region containing the predicted SPBRE site and the promoter region [Lazarevic et al., 1999] was fused to a promoter-less *lacZ* gene. The *P_aimR_* fragment consists of SPBRE site, which is followed by the promoter (Figure 4G). The construct was integrated into the *amyE* locus of the strains TS03 (no *yopR*) and KK101 (*yopR*^G407A^), resulting in the strains KK100 (*P_aimR_-lacZ*) and KK098 (*P_aimR_-lacZ yopR*^G407A^), respectively. The strain KK139 (*P_aimR_-lacZ yopR*) was generated by transformation of KK100 with the plasmid pKK005. Next, we cultivated the bacteria for 2.5 h at 37 °C, shifted the cultures for to 50 °C and continued the cultivation for 2.5 h. The activity of the β-galactosidase was measured every half hour (Figure 4G). As expected, the *P_aimX_* promoter was active over a period of 2.5 h in the strain KK100 lacking *yopR*. The decrease in β-galactosidase activity is probably due to the thermosensitivity of the enzyme. Similarly, the *P_aimX_* promoter in the strain KK098 synthesizing the thermosensitive YopR^G136E^ protein also stayed active within 2.5 h of cultivation. By contrast, β-galactosidase activity in the strain KK139 carrying the wild type *yopR* allele was already greatly reduced after 30 min of incubation. To conclude, β-galactosidase activity assays revealed that YopR is a DNA-binding protein that binds to the *yopJ-aimR* intergenic region of the SPβ genome. We could also confirm that YopR^G136E^ does indeed respond to the ambient temperature *in vivo.* However, the fact that the *P_aimR_* promoter was active at 37 °C when either YopR or YopR^G136E^ was present suggests that other factors exist to inactivate the *P_aimX_* promoter and to prevent entry into the lytic cycle.

### Structural analysis of YopR reveals protein fold typical for tyrosine recombinases

In accordance with our observation that YopR is a DNA binding protein, bioinformatical analysis using the amino acid sequence of YopR identified a potential DNA breaking-re-joining catalytic domain (InterProScan5), and BLAST analysis showed weak similarities to potential integrases and recombinases. Hence, to gain molecular insight into the mode of action of YopR, we performed a crystallographic analysis. After overproduction of YopR fused to a C-terminal His_6_-tag in *E. coli*, the protein was purified using Ni-NTA-affinity and size-exclusion chromatography, concentrated, and used for crystallization screens. Crystals formed within one week of incubation, and we determined the structure of YopR at a resolution of 2.8 Å by molecular replacement using the N-terminal domain (residues 1-105) of the YopR Alphafold model [Jumper et al., 2021] and manually building of the C-terminal part (residues 106-320) (Table 2). Two YopR molecules were present in the crystallographic asymmetric unit (Figure S5). The monomers interact only weakly via residues of mainly two *α*-helices (helix H’ and I) (Figure S5D-F). As YopR elutes as a monomer in analytical SEC experiments (Figure S1A), the formation of a potential YopR dimer appears to be an artifact of crystal packing (Figure 5, and S5). The YopR monomer consists of 15 *α*-helices and 4 β-strands (Figure 5A, B; PDB-ID: 8a0a). The overall structure of a YopR monomer shows a high similarity to bacterial and phage tyrosine recombinases (Figure 5E-F and Figure S6) [Meinke et al., 2016; Grindley et al., 2006]. As other proteins with a tyrosine recombinase fold, YopR harbors a CB (core-binding) domain (Gly1-Lys84, helix A-D) at the N-terminus, a catalytic (CAT) domain at the C-terminus (Tyr108-Ala320, helix F-M), which are connected by a central unfolded linker (Gly85-Leu107, helix/loop E) (Figure 5). While amino acid sequence alignments of YopR to tyrosine recombinases do not yield a high percentage of sequence identity (below 25 percent), structural comparison using PDBefold identifies the highest similarity to the structurally and functionally characterized tyrosine recombinases Cre from *E. coli* phage P1 [Duyne, 2001; Ennifar et al., 2003], XerH from *Helicobacter pylori* [Bebel et al., 2016], and the integrase (Int) from *E. coli* phage lambda [Kwon et al., 1997; Biswas et al., 2005; Aihara et al., 2003] (Figure 5E-H and Figure S6). Most of these tyrosine recombinases have been crystallized in complex with DNA to elucidate the recombination mechanisms in detail, while YopR was crystallized in its apo form. When YopR is structurally aligned to these proteins, the RMSD values for the CB (N-terminal domain) and CAT (C-terminal domain) range between 4 and 6 Å (Table 3). Overlay of YopR with structures of Cre (PDB-ID: 1q3u) [Ennifar et al., 2003], XerH (PDB-ID: 5jk0) [Bebel et al., 2016], and lambda Int (PDB-ID: 1p7d) [Aihara et al., 2003] bound to DNA, revealed the requirement of immense structural rearrangements of YopR during potential DNA binding. While the CB domain in the YopR crystal structure (in absence of DNA) is rotated by approximately 180° vertically and 45° horizontally compared to Cre, XerH and Lambda Int, the CAT domain needs to be flipped by 90° horizontally (Figure 5B, E-H and Figure S6). We hypothesize that the observed organization of YopR might be a crystallographic artefact since YopR was not co-crystallize with DNA. The flexibility for the proposed rearrangements is provided by the approximately 22 residue long linker present between the CB and CAT domain (labelled as helix E’ located between *α*-helices E and F), which is also present in Cre, XerH and Lambda Int (Figure 5 and Figure S6). This structural rearrangement would also generate a positively charged groove between the CB and CAT domains of YopR suitable for DNA interaction (Figure 5C, D).

**Figure 5.**
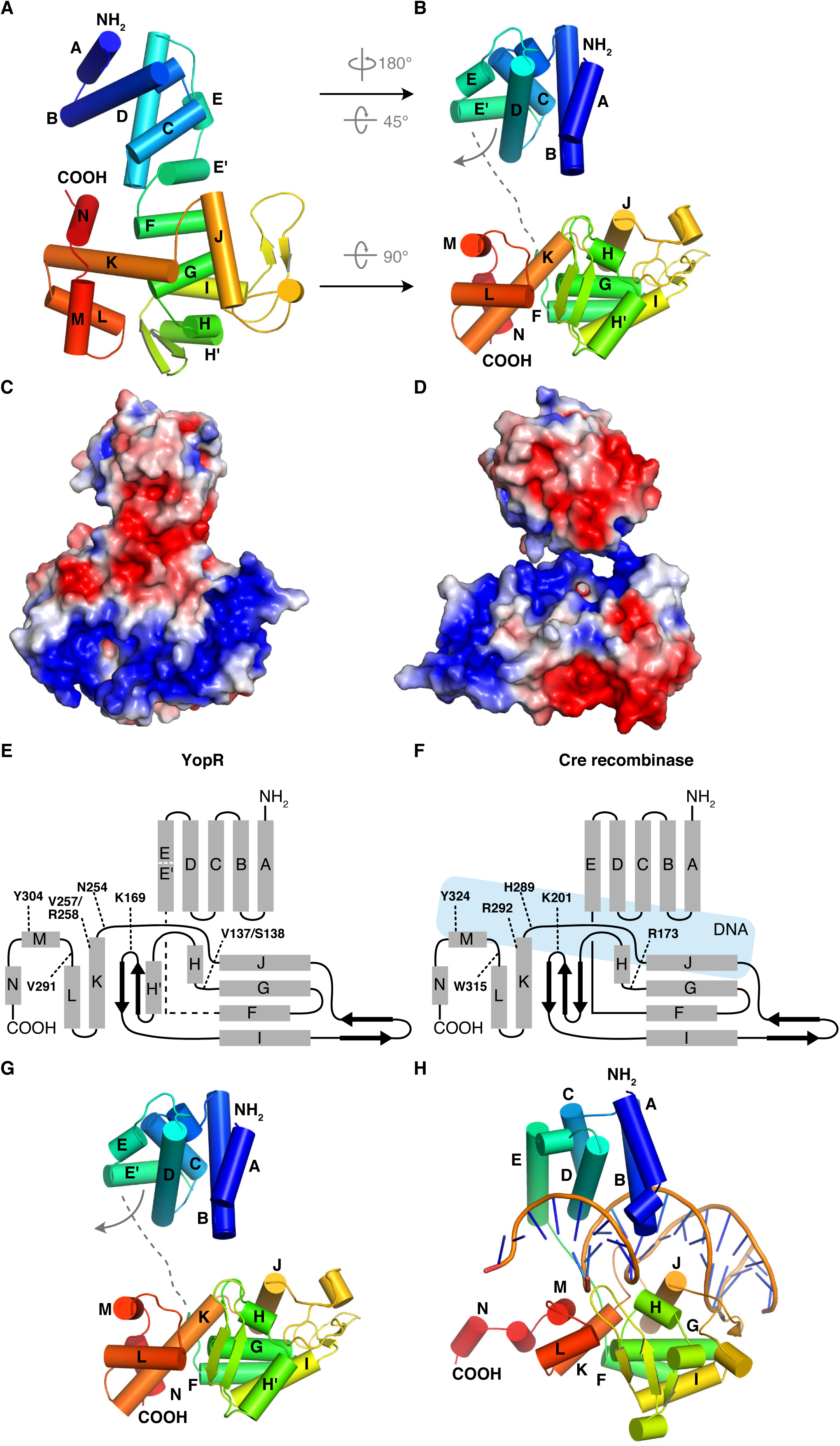
Structural analysis of the YopR wild type protein. **A.** Overall fold of YopR and representation of the 15 alpha-helices and 4 β-strands. **B.** Structural rearrangements of the N- and C-terminal domains of YopR required for potential DNA binding and formation of a negatively charged groove. **C, D.** Electrostatic surface potential of YopR as present in the experimentally determined crystal structure (**C**) and after the proposed structural rearrangements required for DNA binding (**D**). YopR is colored by electrostatic surface potential, as calculated by APBS using the Pymol plug in. The color scale is the same for all proteins, ranging from - 3 to + 3 kT/e, with negative charges in red and positive charges in blue. **E, F.** Schematic overview of YopR (**E**) topology compared to the topology of the well-studied tyrosine recombinase and close structural homolog Cre (**F**). Residues crucial for enzymatic function of tyrosine recombinases are labelled. Location of DNA in Cre is indicated in light blue. **G, H.** Comparison of the overall fold of YopR (**G**) to Cre in its DNA bound state (PDB 1q3u) [Ennifar et al., 2003].

**Table 3.** Comparison crucial residues for the activity of tyrosine recombinases to those found in YopR.

| **YopR**<br>PDB code: 8A0A | **Cre**<br>PDB code: 1Q3U | **XerH**<br>PDB code: 5JK0 | **Lambda Int**<br>PDB code: 1P7D |
| --- | --- | --- | --- |
| RMSD (Å) N- / C-terminus | 4.61 / 4.53 | 6.04 / 5.14 | 4.03 / 4.06 |
| V137/S138 | R173 | R213 | R212 |
| E144 | E176 | E216 | D215 |
| K169 | K201 | K239 | K235 |
| N254 | H289 | H309 | H308 |
| V257/R258 | R292 | R312 | R311 |
| V291 | W315 | H335 | H333 |
| Y304 | Y324 | Y344 | Y342 |
| References | [Ennifar et al., 2003;<br>Van Duyne, 2015;<br>Meinke et al., 2016] | [Bebel et al.,<br>2016] | [Aihara et al.,<br>2003; Biswas et al.,<br>2005; Landy, 2015] |

### Perturbations in secondary and higher order structure of YopR^G136E^

The difference in melting temperature suggested to us that YopR^G136E^ may differ in its conformation from YopR causative for the higher instability of the former. We thus employed hydrogen/deuterium exchange mass spectrometry (HDX-MS) to ascertain potential differences in conformation between both YopR variants. HDX-MS probes the rate with which the amide protons contained in the peptide bond exchange for protons from the solvent, the degree of which is an intrinsic characteristic of an amino acid sequence, whether the amide protons establish hydrogen bonding interactions and their solvent accessibility. Incubation of a protein in D_2_O-containing solvent visualizes this exchange rate because a successful exchange results in a mass shift that can be quantitated by MS. Hence, HDX-MS may provide insights into a protein’s higher order structure and conformational dynamics [Masson, 2019]. The comparison of the HDX of YopR and YopR^G136E^ reveals that the latter incorporates more deuterium in many parts of the protein (Figure 6A and Figure S7). These regions are mainly located in the CAT domain, but less extensive changes are also apparent in the CB domain (e.g., residues 51-63, Figure 6B). While the G136E site of variation cannot be directly compared in this way because different peptides are obtained for either protein, areas in spatial proximity of the site of variation similarly show elevated HDX for YopR^G136E^ (residues 250-258, Figure 6B). Illustration of all the YopR areas in which YopR^G136E^ incorporates more deuterium on its crystal structure highlights that almost the entire CAT domain and the proximal portions of the CB domain (Figure 6C) are affected. As the HDX rate is inversely correlated with the degree of higher order structure, the higher HDX of YopR^G136E^ than native YopR is indicative for a less ordered conformation in most parts of this protein variant coinciding with its lower melting temperature (Figure 3B).

**Figure 6.**
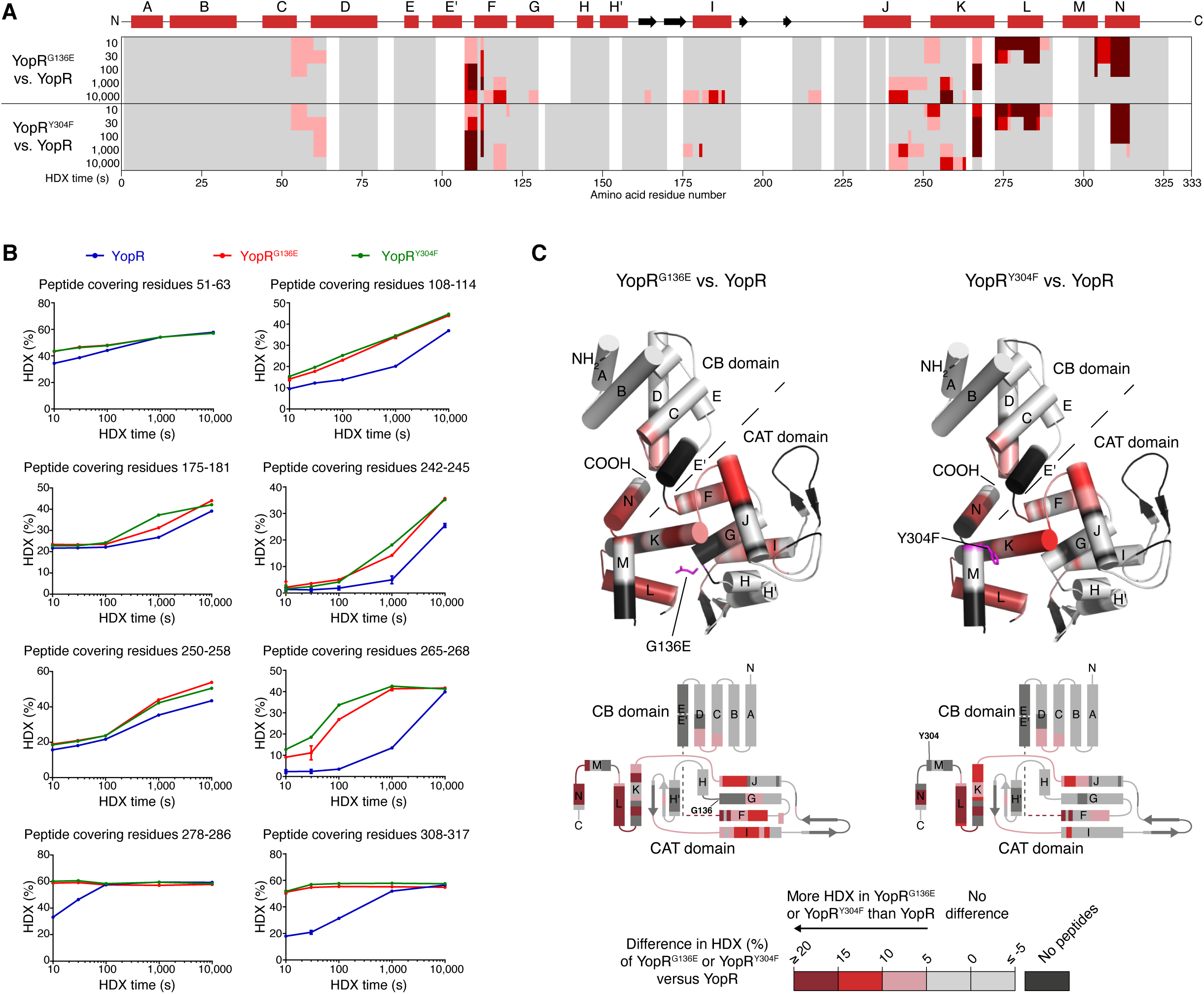
YopR^G136E^ and YopR^Y304F^ exhibit perturbations in higher order structure compared to YopR. **A.** The difference in HDX-MS between YopR^G136E^ and YopR or YopR^Y304F^ and YopR is displayed on the amino acid sequence of YopR. Different tones of red indicate residues that incorporate more deuterium in the YopR^G136E^ and YopR^Y304F^ variants than in native YopR indicating perturbations in their higher order structure. The secondary structure of YopR is schematically illustrated above. **B.** Representative peptides of YopR exhibiting differences in HDX between YopR (blue trace), YopR^G136E^ (red trace) and YopR^Y304F^ (green trace). Data represent the mean ± s.d. of n=3 replicates. **C.** The difference in HDX-MS between YopR^G136E^ or YopR^Y304F^ and YopR (see panel A) was mapped onto the YopR crystal structure. Hereby, each residue was color-coded as per the highest difference in HDX observed at any time-point. Red colors represent higher HDX in YopR variants and grey color no HDX difference compared to native YopR. YopR residues for which no peptides could be identified in HDX-MS are colored black. The G136E and Y304F sites of variation are shown as sticks colored in magenta.

### Mutational and functional analysis of YopR

Tyrosine recombinases are site-specific DNA modifying enzymes that bind, cleave, strand exchange, and rejoin DNA at their respective, typically palindromic, recognition target sites. The function of this enzyme family has been best studied in detail for the Cre, Xer and Lambda Int recombinases. For each of these recombinases, detailed molecular studies have led to the identification of residues that are crucial for DNA recombination activity [Grindley et al., 2006; Ennifar et al., 2003; Aihara et al., 2003; Bebel et al., 2016; Jayaram et al., 2015]. The analyzed residues listed in Table 3 resemble a core active site of tyrosine recombinases, with highly conserved catalytic core: R, D/E, K, H, R, H/W and Y. These residues engage the scissile phosphate and a tyrosine nucleophile that forms a 3ʹ phosphotyrosine linkage with the cleaved DNA. When exemplarily compared to the closest structural homologs Cre, XerH, and Lambda Int, only three of the seven residues crucial for catalysis are present in YopR (Table 3, Figure 5 D-E, Figure 7 and Figure S8). Especially the crucial arginine and histidine residues are substituted by serine, valine and asparagine in YopR (Table 3, Figure 5, Figure 7, Figure S8). The first conserved residue also present in YopR is K169, which is homologous to K201 in Cre, K239 in XerH and K235 in Int (Table 3, Figure 7, Figure S8). In Cre recombinase, K201 contacts a base adjacent to the cleaved phosphate in the minor groove of the DNA and the residue is positioned in a flexible loop. Alanine mutants of this lysine residue in XerD and Cre were defective in recombination [Duyne et al., 2001]. The second conserved residue is a nucleophilic tyrosine (Y304 in YopR, Y324 in Cre, Y344 in XerH, and Y342 in Int; Table 3, Figure 7, Figure S8) that is described to form a phosphor-protein covalent linkage. As shown for Cre and Lambda Int, it attacks the scissile phosphate forming a 3ʹ-phosphotyrosine linkage and resulting in a free 5ʹ-HO during DNA cleavage [Duyne et al., 2001; Biswas et al., 2005]. The third conserved residue also found in YopR is E144. In some tyrosine recombinases such as Lambda Int it is substituted by an aspartate (Table 3). This residue appears to contribute indirectly to transition state stabilization by hydrogen bonding to catalytic residues and promoting the integrity of the active site [van Duyne et al., 2001; Biswas et al., 2005; Meinke et al. 2016]. The other residues (histidines and arginines) required for catalytic activity, synapsis formation and stabilization of the transition state during recombination are not conserved in YopR and are replaced by serine, valine and asparagine residues (Table 3, Figure 7, Figure S8). Hence, it is questionable if YopR also functions as a DNA-cleaving recombinase or rather has lost this enzymatic activity.

**Figure 7.**
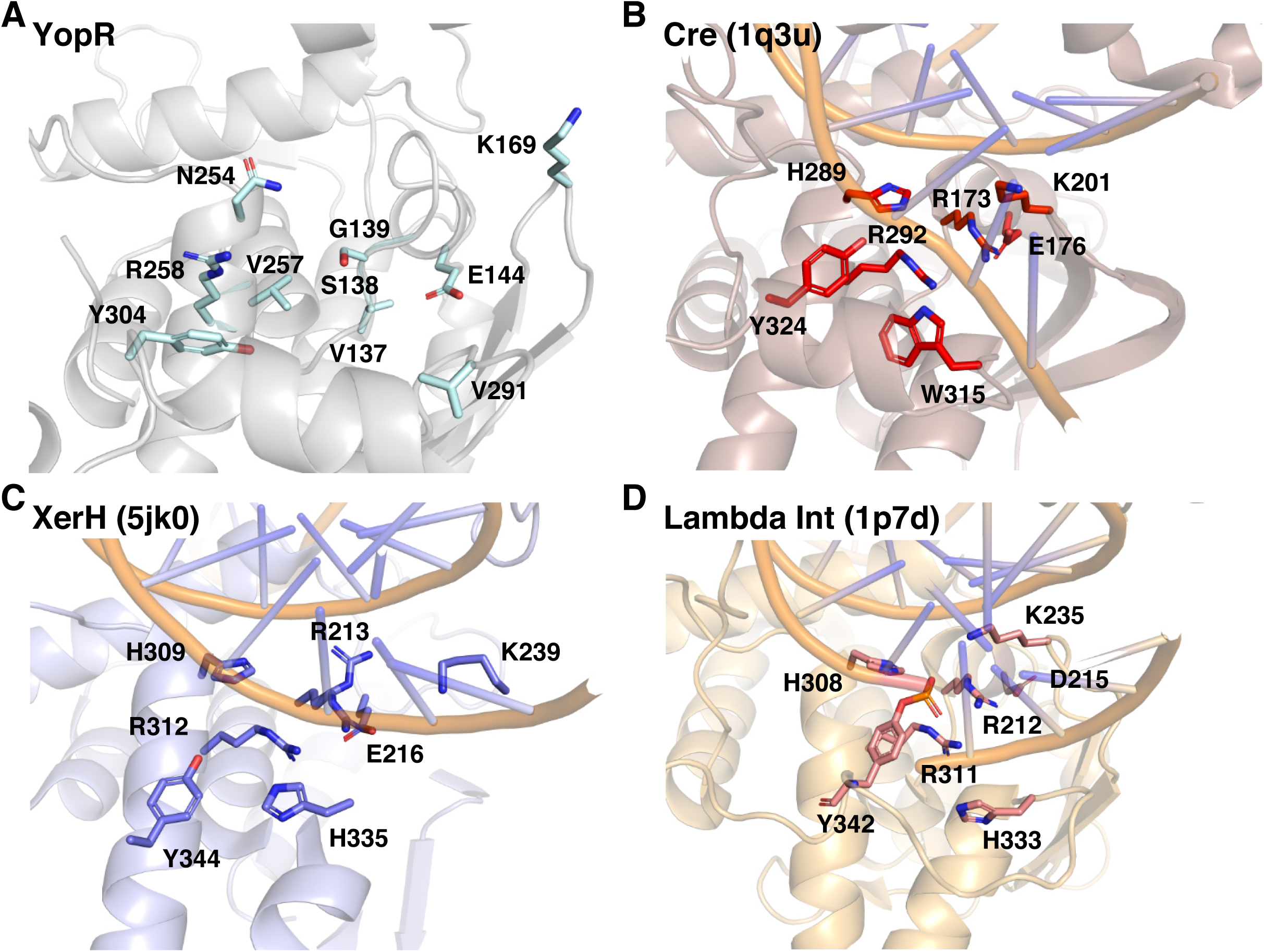
Detailed structural comparison of YopR and the core catalytic residues of tyrosine recombinases. **A.** Zoom into the potential DNA binding region and catalytic site of YopR. Residues that were compared to crucial catalytic residues of tyrosine recombinases in Table 3 are shown as light blue sticks and labelled. **B-D.** Comparison of the core catalytic residues in the tyrosine recombinases Cre (PDB 1q3u; Ennifar et al., 2003) (**B**), XerH (PDB 5jk0; Bebel et al., 2016) (**C**) and Lambda Int (PDB 1p7d; Aihara et al., 2003) (**D**). The catalytic residues are shown as red, blue and pink sticks, respectively. Detailed overlay of each residue with the corresponding amino acid residue found in the structure of YopR are shown in Supplementary Figure S7.

It has previously been shown that replacement of the nucleophilic tyrosine in the tyrosine recombinases Cre, XerH and Lambda Int by either a phenylalanine or alanine residue renders the recombinase inactive [Grindley et al., 2006; Ennifar et al., 2003; Aihara et al., 2003; Bebel et al., 2016; Jayaram et al., 2015]. Since this potential nucleophilic tyrosine (Y304 in YopR) is also found in YopR and to further investigate the biological function of YopR, we introduced point mutations in the *yopR* gene leading to a replacement of Y304 by alanine or phenylalanine. We further generated a YopR mutant in which the conserved catalytical residue K169 is exchanged by an alanine. Each of these proteins could be produced heterologously in *E. coli* and purified (Figure S1G). We first characterized these YopR variants biochemically and analyzed their folding and oligomeric state via analytical size exclusion. All tested variants except YopR ^Y304A^ showed a monomeric protein peak in this analysis (Figure S1D-G). YopR ^Y304A^ possessed a peak shoulder, showed bands of degradation in SDS-PAGE analysis and large parts of the protein were insoluble during overproduction indicative for problems in protein folding and stability (Figure S1E, H). We further tested the thermostability of these variants using nanoDSF. While the YopR^K169A^ variant showed the same thermal unfolding behavior as the wild type YopR protein, the thermal stabilities of the YopR^Y304F^ and YopR^Y304A^ variants were reduced by about 6 °C (Figure 3B) comparable to the initially tested heat sensitive YopR^G136E^ variant. This inference is also substantiated by changes in HDX-MS of YopR^Y304F^ compared to native YopR, which are highly reminiscent in quality and quantity to those observed for the heat sensitive YopR^G136E^ variant (Figure 6).

Next, we assessed the YopR^K169A^ and YopR^Y304F^ variants for their *in vitro* DNA binding affinity via ITC using the same regions as described previously. While the YopR^K169A^ variant possessed a DNA binding affinity to the fragment upstream of *aimR* in a similar range as YopR wild type in the ITC analysis (0.238 ± 0.08 µM for YopR wild type and 0.346 ± 0.15 µM for YopR^K169A^), the Y304F exchange resulted in a phenotype comparable to the heat sensitive G136E mutant (1.69 ± 0.69 µM for YopR^G136E^ and 1.34 ± 0.5 µM for YopR^Y304F^) (Figure S3). To further consolidate this notion, we performed HDX-MS on YopR, YopR^G136EF^ and YopR^Y304F^ in presence of DNA and compared the HDX rate of those to their respective apo states (Figure 8). The protein regions exhibiting reduced HDX in presence of DNA are similar for all three variants except for the linker region between helices E and F, and the C-terminal YopR portion encompassing helices K to N (Figure 8A). These concur with the areas where YopR^G136E^ and YopR^Y304F^ showed the most pronounced defects in higher-order structure (Figure 6). Collectively, these data suggest that both variants are, in principle, still capable of DNA coordination via the CAT domains’ N-terminal portion and that their reduced DNA binding affinity is causally linked to structural defects in the C-terminal part of the CAT domain (Figure 8C).

**Figure 8.**
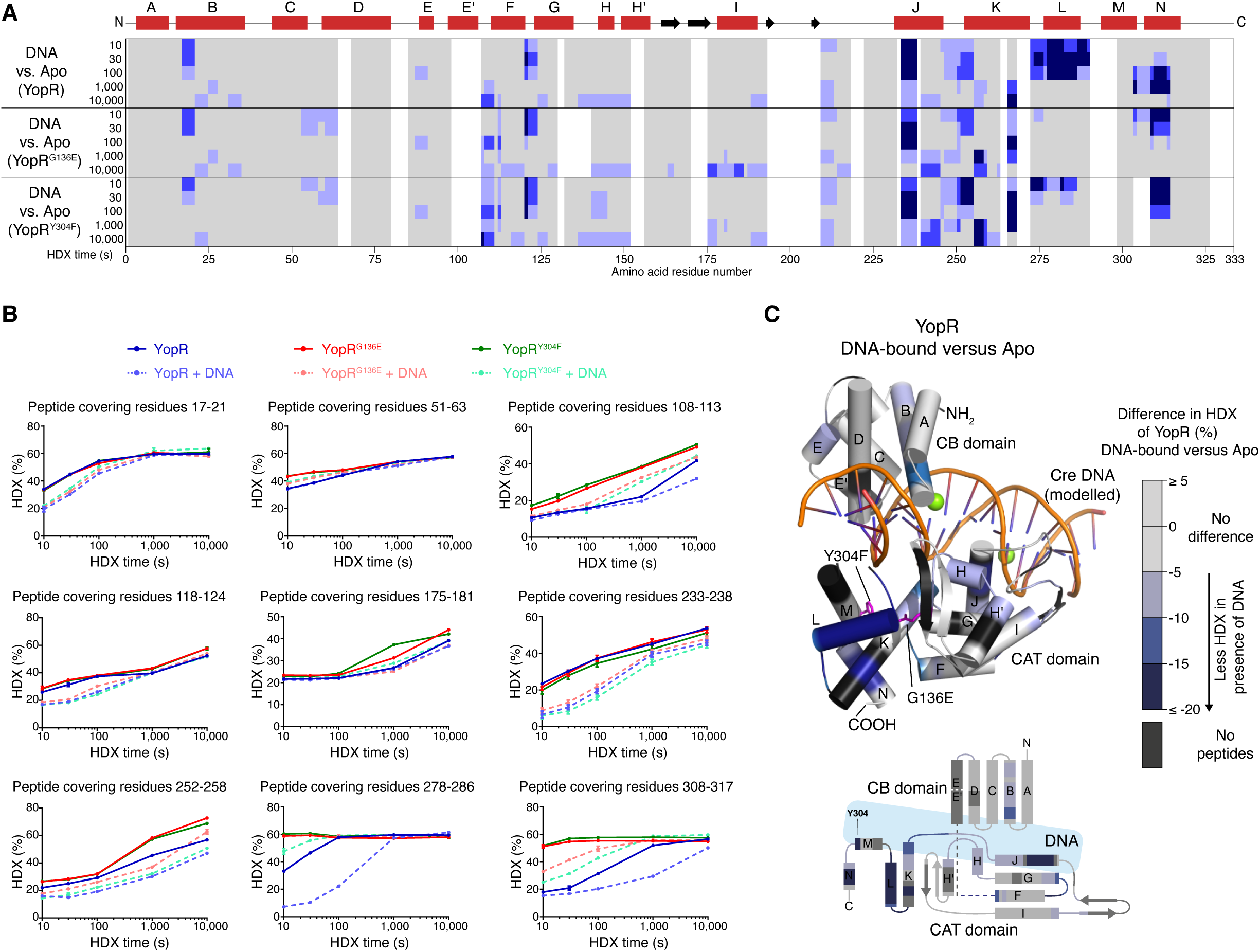
DNA binding of YopR, YopR^G136E^ and YopR^Y304F^ by HDX-MS. **A.** The difference in HDX-MS between the DNA-bound and apo states of YopR, YopR^G136E^ and YopR^Y304F^ is displayed on the amino acid sequence of YopR. Different tones of blue indicate reduced deuterium incorporation in presence of DNA. The secondary structure of YopR is schematically illustrated above. **B.** Representative peptides of YopR and variants thereof exhibiting differences in HDX with respect to DNA. Data represent the mean ± s.d. of n=3 replicates. **C.** The difference in HDX-MS between the DNA-bound and apo state of native YopR was projected onto a model of DNA-bound YopR. The model was generated by superimposing the CAT domains of YopR (residues 107-318) and DNA-bound Cre (PDB 1q3u [Ennifar et al.], 2003; residues 130-341) followed by superposition of the YopR CB domain (residues 1-106) onto that of DNA-bound Cre (residues 10-129). YopR residues were color-coded as per the highest difference in HDX observed at any time-point. The G136E and Y304F sites of variation are shown as sticks colored in magenta.

### YopR does not show recombination activity

Since YopR possesses a similar fold as tyrosine recombinases and at least harbors the catalytically important tyrosine and lysine residue, we wondered if YopR can function as a recombinase *in vitro* using the *attL* and *attR* sites of the SPβ phage [Abe et al., 2017]. To assess the activity of YopR WT, and the YopR^G136E^, YopR^Y304F^ and YopR^K169A^, we employed an *in vitro* recombination assay previously described by Abe et al. with minor changes as we used fluorescently labeled fragments of the *attL* (242 bp, labelled with Cy5) and *attR* (616 bp, labelled with Cy3). If YopR would be able to cleave, strand exchange, and rejoin the SPβ recombination sites, we would expect the appearance of a hybrid DNA fragment with the size of 426 bp carrying both fluorescence labels (Figure S9A). YopR wild type and the tested YopR^G136E^, YopR^Y304F^, and YopR^K169A^ variants were incubated with the labeled *attL* and *attR* sites for 60 min at 37 °C, boiled to release YopR from the DNA and analyzed on a native polyacrylamide gel. Neither YopR wild type, nor any of the tested YopR variants resulted in the formation of a hybrid DNA fragment carrying both fluorescence labels, which would be indicative for a DNA recombination event of the SPβ *att* sites (Figure S9B).

Next, we assessed the *in vivo* activities of the YopR^K169A^, YopR^Y304F^ and YopR^Y304A^ variants by performing a heat shock experiment. For this purpose, we introduced the *yopR*^A911T^ (Y304F)*, yopR*^TAT910-912GCG^ (Y304A) and *yopR*^AAG505-507GCG^ (K169A) alleles into the *amyE* locus of the strain KK002 (SPβ c2). Next, we deleted the native *yopR*^G407A^ copy from the SPβ c2 genome resulting in the strains KK120 (SPβ c2 [*yopR*^G407A^*::ermC*] *P_alf4_-yopR*^A911T^), KK121 (SPβ c2 [*yopR*^G407A^*::ermC*] *P_alf4_-yopR*^TAT910-912GCG^) and KK138 (SPβ c2 [*yopR*^G407A^*::ermC*] *P_alf4_-yopR*^AAG505-507GCG^). The fact that it was possible to delete the native *yopR*^G407A^ allele indicates that the YopR^K169A^, YopR^Y304F^ and YopR^Y304A^ variants were functional because they prevented the entry of the SPβ *yopR* mutant to enter the lytic cycle under unstressed conditions (Figure 3C). Consistent with the nanoDSF experiments (Figure 3B), only the YopR^Y304F^ and YopR^Y304A^ variants but not YopR^K169A^ showed increased temperature sensitivities (Figure 3C). Thus, the residues G136 and Y304 are critical for the thermal stability of YopR. Moreover, it is unlikely that YopR functions as a recombinase because the replacement of the tyrosine residue, which is crucial for enzyme catalysis of known recombinases (see above), still allows the phage to enter the lytic cycle.

### Heat induction of the lytic cycle of SPβ c2 does not depend on SprB

Previously, it was shown that SprA and SprB are required for SPβ prophage excision during sporulation (Figure 1) [Abe et al., 2014]. We were wondering if the heat-inducible SPβ excision of the c2 mutant also depends on SprA and SprB. To test this idea, we deleted the *sprA* and *sprB* genes in the strain KK002 carrying SPβ c2. Genome sequencing verified the deletion of the *sprA* and *sprB* genes in the strains KK124 (*sprA::aphA3*) and KK125 (*sprB::aphA3*), respectively. Both strains also carried the four mutations that were identified in the SPβ c2 phage (Table 1). Next, we performed a heat shock experiment and assessed cell lysis of the strains KK124 (SPβ c2 [*sprA::aphA3*]) and KK125 (SPβ c2 [*sprB::aphA3*]). As shown in Figure 9, both strains still lysed in the absence of either SprA or SprB. However, only the *sprB* mutant produced infectious SPβ c2 phage particles as verified by plaque assays using the prophage free strain TS01 (data not shown). We conclude that the heat-inducible SPβ excision of the c2 mutant also depends on the serine recombinase SprA while the heat-inducible induction of lysis is independent of SprA and SprB.

**Figure 9.**
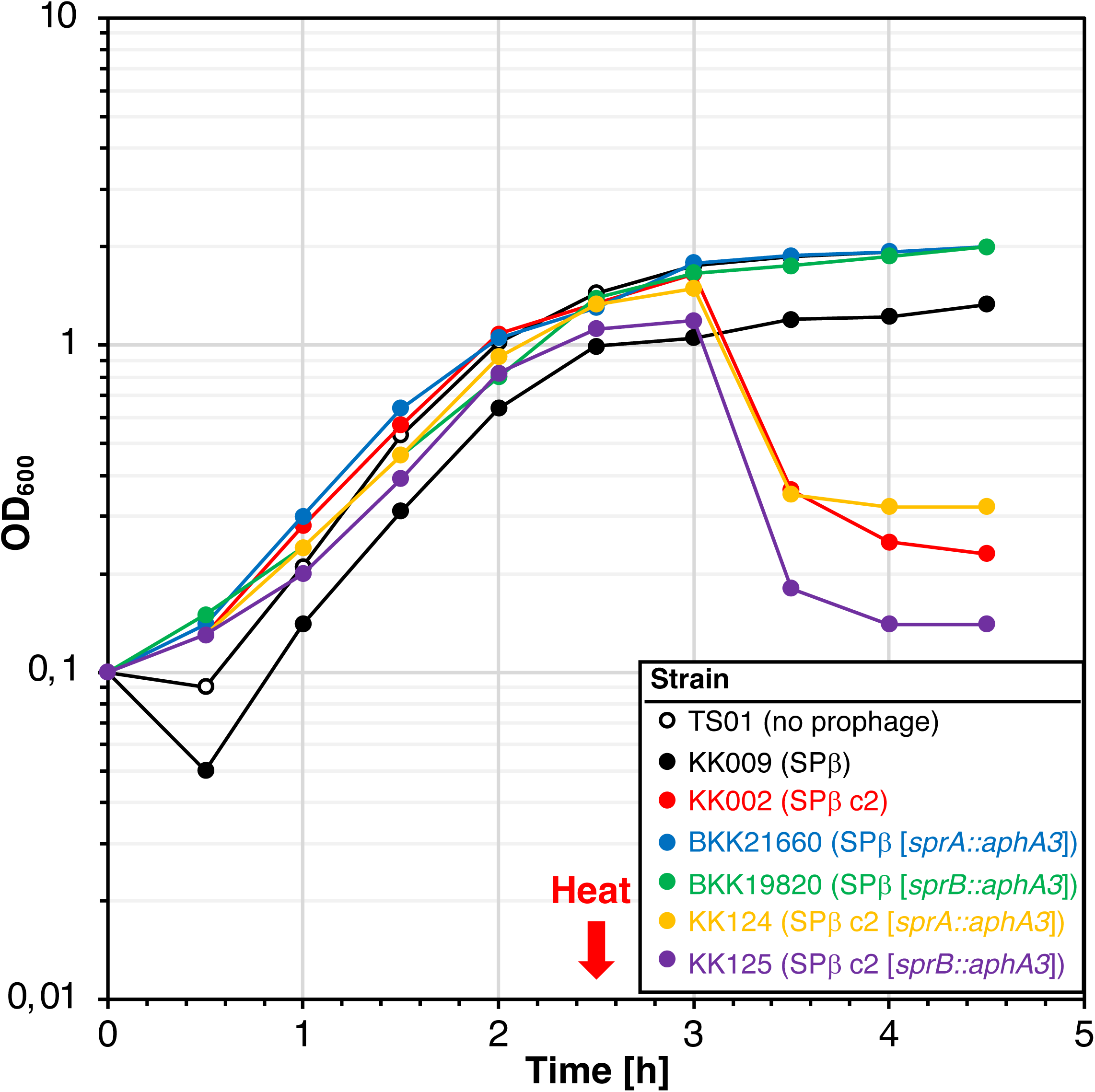
Heat-induced cell lysis of *B. subtilis* carrying SPβ c2 does not require SprA or SprB. The indicated strains were grown for 2.5 h at 30 °C, heat shocked at 50 °C for 10 min and further cultivated at 30 °C. The square brackets indicate the *sprA* and *sprB* deletions in the phage genome. The OD_600_ was monitored over time.

### Identification of YosL, a novel component of the lysis-lysogeny management system of SPβ

When performing heat shock experiments with the strain KK001 carrying the SPβ c2 prophage, we observed that approximately 10% of the cells survived. Characterization of individual isolates of the surviving cells revealed that the lytic cycle of SPβ c2 was still inducible by heat, indicating the presence of intact prophages (data not shown). We speculated that among the surviving cells, there could be SPβ c2 mutant variants that had acquired intra- or extragenic mutations, which would prevent heat inducibility of the lysogens. To isolate SPβ c2 mutant variants that had lost the heat-sensitive phenotype, a small-scale adaptive laboratory evolution experiment was performed. For this purpose, the strain KK001 (SPβ c2) was grown at 37 °C until the culture reached an OD_600_ of about 0.8, heat shocked for 10 minutes at 50 °C and incubated for additional 6 hours at 37 °C. After 2 hours of incubation cell lysis took place, which could be recognized by the decrease in optical density. When the culture again reached an OD_600_ of 0.8, fresh medium was inoculated to an OD_600_ of 0.1 and incubated overnight at 37 °C. Next day, cells from the overnight culture were used to inoculate fresh medium to an OD_600_ of 0.1 and the heat shock cycle was repeated. After performing this cycle for 7 days, cell lysis could no longer be induced by heat, suggesting that genes of the SPβ lysis-lysogeny management system had probably acquired detrimental mutations. Next, 15 clones from the evolved culture were isolated, and the presence of the prophage was verified by a sublancin assay. The culture supernatants of five randomly selected clones were used to generate lysogens in the background of the prophage-free strain TS01. In two of the 19 isolated prophage-containing strains that were designated as KK026 and KK027, cell lysis could no longer be induced by heat (Figure 10A). Genome and Sanger sequencing revealed that the SPβ c2 prophage had acquired mutations in the *yosL* gene in both strains. In strain KK026, a single-nucleotide insertion (+A1) would lead to a frameshift that truncates the YosL protein after only four of the predicted 117 amino acids. The strain KK027 had a single-nucleotide exchange in *yosL* (T125A) that likewise would truncate YosL after 59 amino acids. We also observed that cell lysis of the strains KK026 (SPβ c2 [*yosL*^+A1^]) and KK027 (SPβ c2 [*yosL*^T125A^]) could not be induced by treatment with mitomycin C (Figure 10B). Thus, YosL is essential for the heat- and mitomycin C-dependent induction of the lytic cycle of SPβ c2.

**Figure 10.**
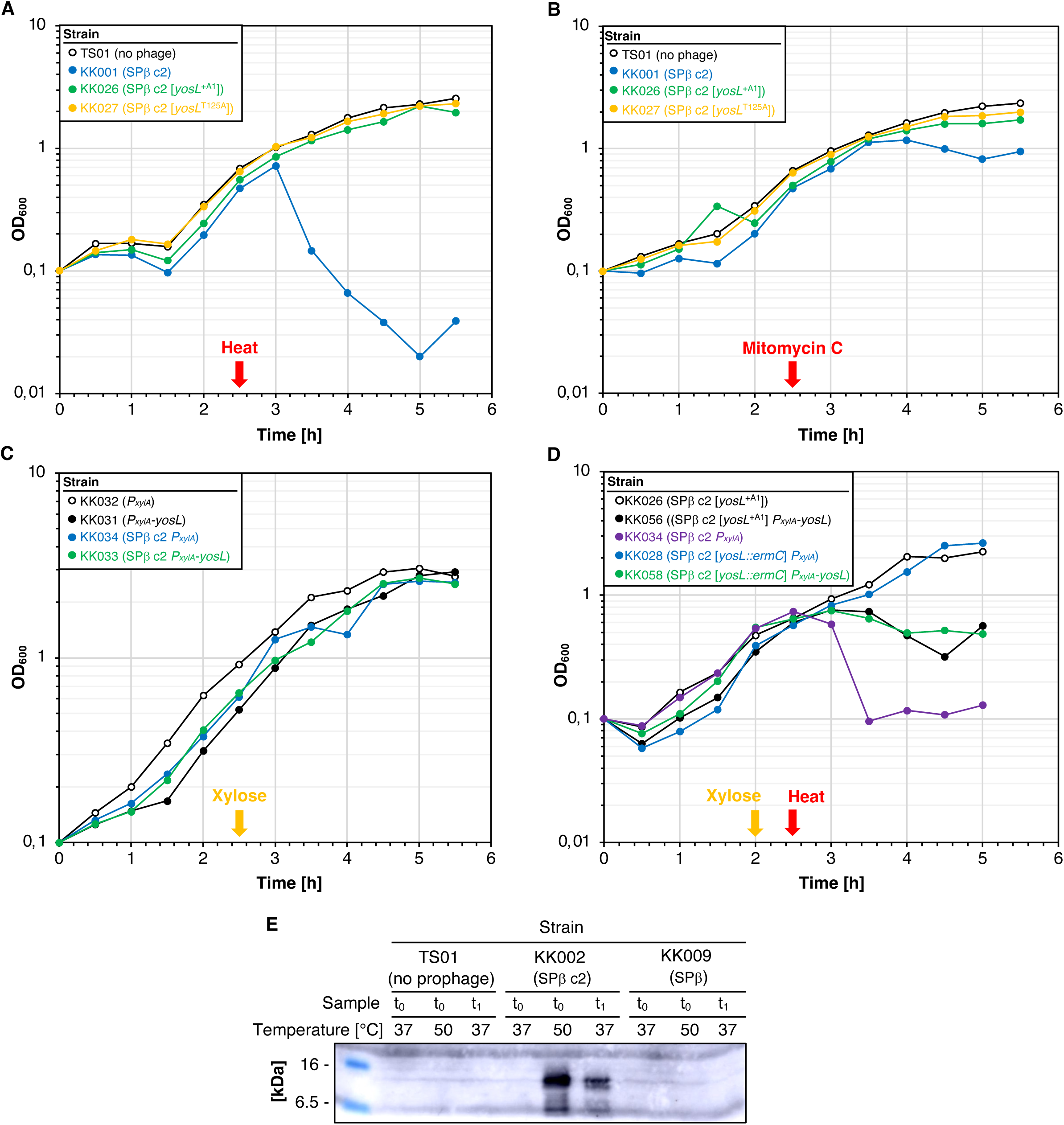
YosL, a novel player in the lysis/lysogeny management system of SPβ. **A.** Characterization of evolved SPβ c2 suppressor mutants. All strains were grown for 2.5 h at 37 °C, heat shocked at 50 °C for 10 min and further cultivated at 37 °C. The OD_600_ was monitored over time. **B.** Effect of mitomycin C on prophage induction in strains carrying SPβ c2 and its suppressors during growth in LB at 37 °C. Mitomycin was added to a final concentration of 0.5 µg ml^-1^ and OD_600_ was monitored over time. **C**. Effect of xylose (0.5% (w/v)) on prophage induction in the indicated strains that were grown in LB. **D.** Effect of xylose (0.5% (w/v)) and heat (10 min at 50 °C) on prophage induction in the indicated strains that were grown in LB. **E.** Cellular amounts of YosL were determined by Western blotting. Crude extracts of the indicated strains were isolated from exponentially growing LB cultures at timepoints t_0_ (before the heat shock), t_1_ (immediately after the heat shock) and t_2_ (30 min after the heat shock). After electrophoresis in a 15% SDS-PAGE and transfer onto a PVDF membrane, YosL was detected by rabbit polyclonal antibodies raised against YosL-His_6_. 20 mg of crude extracts were applied.

Since the *yosL* gene is not expressed in the dormant state of the SPβ prophage [Kohm et al., 2022], *yosL* transcription must be activated when the phage enters the lytic cycle in a YopR-dependent manner. However, to assess whether YosL can trigger the lytic cycle independent of YopR G136E, we constructed the plasmid pRH168 allowing the xylose-dependent expression of the *yosL* gene from the *ganA* locus. YosL is not toxic for the bacteria because the prophage-free control strains KK31 (*P_xylA_-yosL*) and KK32 (*P_xylA_,* control) grew in the presence of xylose (Figure 10C). Moreover, the expression of *yosL* in the strain KK033 (SPβ c2 *P_xylA_-yosL*) carrying the prophage did not result in cell lysis (Figure 10C). Next, we tested whether the artificially expressed *yosL* gene complements the spontaneously inactivated *yosL*^+A1^ allele and the deletion of *yosL* in the strains KK056 (SPβ c2 [*yosL*^+A1^] *P_xylA_-yosL*) and KK058 (SPβ c2 [*yosL::ermC*] *P_xylA_-yosL*), respectively, upon heat shock. The parental strains KK028 (SPβ c2 [*yosL::ermC*]) and KK026 (SPβ c2 [*yosL*^+A1^]) served as controls. As shown in Figure 10D, albeit to a lesser extent compared to the control strain KK034 (SPβ c2 *P_xylA_*), the xylose-dependent expression of *yosL* in the strains KK056 (SPβ c2 [*yosL*^+A1^] *P_xylA_-yosL*) and KK058 (SPβ c2 [*yosL::ermC*] *P_xylA_-yosL*) induced lysis after heat treatment. Thus, YosL depends on YopR^G136E^ to activate the lytic cycle of SPβ.

To assess whether the heat shock affects the synthesis of the YosL protein in the strain KK002 (SPβ c2), we performed a Western blot using polyclonal antibodies raised against YosL. The prophage free strain TS01 and the strain KK009 (SPβ) served as controls. Samples for the preparation of cell-free crude extracts were taken during cultivation at 37 °C before and 30 minutes after applying a heat shock at 50 °C. As expected, YosL was not synthesized in the strains TS01 (no prophage) and KK009 (SPβ) (Figure 10E). By contrast, immediately after the heat shock, the strain KK002 (SPβ c2) synthesized a protein corresponding to YosL (13 kDa). To conclude, we identified YosL as a novel component of the lysis-lysogeny management system that is required for the heat- and mitomycin C-dependent induction of the lytic cycle of the SPβ c2 mutant.

## DISCUSSION

Previously, it has been shown that the *yopR* gene codes for the master repressor of the lytic cycle of the SPβ prophage, which is present in the *B. subtilis* laboratory strain 168 [Brady et al., 2021; Kohm et al., 2022]. Here, we confirmed that YopR is a key player in the lysis-lysogeny management system of SPβ. We show that a single mutation in the *yopR* gene enables induction of the SPβ c2 lytic cycle by transiently increasing the cultivation temperature. This finding suggested that either the stability or the folding of the encoded YopR^G136E^ protein is influenced by the ambient temperature of the bacterial culture. The first possibility could be excluded because the cellular levels of YopR^G136E^ were not affected by the ambient temperature (Figure 3A). By contrast, biochemical characterization of the YopR^G139E^ variant revealed that its unfolding temperature was decreased by about 10 °C and that this variant exhibited less higher order structure (Figure 3B and Figure 6). Thus, a single amino acid exchange in the YopR protein rather renders the folding of the protein sensitive to temperature. Our structural characterization of YopR revealed that the monomer shows a high similarity to bacterial and phage tyrosine recombinases and that the Y304 residue might be crucial for the DNA-binding activity of the protein (Figure 6 and S6). It is interesting to note that the replacement of tyrosine by phenylalanine or alanine at position 304 also makes the YopR^Y304F^ and YopR^Y304A^ variants temperature sensitive, albeit to a lesser extent than the G136E exchange in YopR^G136E^ (Figure 3B and 3C). However, the precise role of Y304 must be determined by structural characterization of YopR-DNA complexes. Like for many other systems, temperature-sensitive variants of DNA binding proteins, including the repressor of the lysogenic phage *λ* were shown to be helpful to study the underlying molecular mechanisms by which DNA-binding transcription factors exert their function [Lieb, 1996, 1981; Wissmann et al., 1991; Hurme et al., 1996]. Moreover, the entry into the host environment and the accompanied temperature change serve as cues for thermosensing transcription factors to control virulence gene expression in pathogenic bacteria [Hurme et al., 1997]. Temperature-sensitive DNA binding transcription factors have also been employed for constructing expression systems that can be triggered by an ambient temperature [Jechlinger et al., 2005; Valdez-Cruz et al., 2010]. However, it is rather unlikely that the temperature serves as a signal for the SPβ wild-type prophage to enter the lytic cycle due inactivation of YopR because a heat shock does affect its repressor function. Recently, it has been suggested that the transcription factor AimR might be required to relieve YopR repression, thereby promoting the entry of SPβ into the lytic cycle [Brady et al., 2021]. It would be interesting to test whether the AimX nc RNA plays a role in this process. However, the environmental or host-derived signal that triggers the inactivation of YopR remains to be identified.

As described above, the overall structure of a YopR monomer shows a high similarity to bacterial and phage tyrosine recombinases (Table 3, Figure 5, Figure S6). However, our analysis indicates that YopR lacks recombinase activity, at least under the tested *in vitro* conditions (Figure S9). Furthermore, the exchange of Y304 to a phenylalanine or alanine residue, facilitated heat induction of the lytic cycle *in vivo* like the G136E variation (Figure 3C). As similar exchanges of the catalytic tyrosine residue in other tyrosine recombinases such as Cre, XerH or Lambda Int render the proteins inactive/incapable of DNA recombination [Grindley et al., 2006; Ennifar et al., 2003; Aihara et al., 2003; Bebel et al., 2016; Jayaram et al., 2015], this may be regarded as an indication that YopR does not act as a tyrosine recombinase despite the structural similarities. Instead, YopR might rather have evolved to a DNA-binding repressor protein. Along this line, YopR might be an evolutionary intermediate that is structurally related to a tyrosine recombinase as well as functionally to regulatory proteins. There are several examples for the evolution of an enzyme to a DNA-binding regulator [Commichau and Stülke, 2015]. For instance, the DNA-binding repressor protein Mlc from *Thermus thermophilus* responds to glucose that binds to a motif, which is conserved in glucose kinases [Titgemeyer et al., 1994; Chevance et al., 2006]. In fact, integrases have been described as acting as repressors to modulate their own synthesis [Piazolla et al., 2006; Chitto et al., 2020]. Probably, the recruitment of tyrosine recombinases as DNA-binding transcription factors is more widespread than previously anticipated.

Previously, it has been shown that SprA and SprB are key components required for the excision of the SPβ genome during sporulation (Figure 1) [Abe et al., 2014]. Here, we could show that lysis of a strain carrying the SPβ c2 allele was induced in when either *sprA* or *sprB* were absent (Figure 4G). The fact that only the strain lacking the *sprA* gene did not produce infective phage particles indicates that the heat-inducible SPβ excision of the c2 mutant also depends on the serine recombinase SprA but probably not on SprB. However, it is not unlikely that additional factors and regulatory circuits control prophage excision during sporulation or due to mitomycin C treatment. First, SprB overproduction was shown to trigger prophage excision but not the lytic cycle [Abe et al., 2014]. Second, a genome-reduced variant of SPβ containing only the *sprA* and *sprB* genes retained the ability to rearrange the *spsM* gene but did not undergo excision during mitomycin C treatment [Abe et al., 2014]. It will be interesting to elucidate whether the yet unknown prophage-derived or the host factors are involved excision of the SPβ.

As mentioned above, temperature-sensitive variants of DNA-binding proteins were helpful to study the functions of DNA-binding transcription factors. Here, we took advantage of the temperature-sensitive YopR^G136E^ variant to identify the YosL protein as a novel component of the lysis-lysogeny management system of SPβ. We showed that the YosL protein was synthesized when a *B. subtilis* lysogen carrying the SPβ c2 prophage was exposed to heat (Figure 10). A bioinformatic analysis revealed that YosL is predicted to possess an Arc-type ribbon-helix-helix domain, which is also present in the structurally homologous repressors Mnt and Arc of the phage P22 that employs both repressors for maintaining the lysogenic state [Burgering et al., 1994; Anderson and Sauer, 2003]. There are two possibilities how the YosL protein might activate the SPβ c2 lytic cycle. First, elevated cellular amounts of YosL could inhibit the synthesis of another factor that prevents the synthesis of components activating the lytic cycle of SPβ c2. Second, YosL might directly activate yet unknown factors triggering the lytic cycle of SPβ c2. However, future structural and functional analysis is required to gain deeper insights into the regulatory role of YosL. Surprisingly, we found that the overexpression of YosL alone did not activate the lytic cycle of SPβ (Figure 10). A similar observation has been made when the transcription factor AimR that activates the expression of the nc RNA AimX, which in turn activates the lytic cycle of SPβ, was overexpressed [Brady et al., 2021]. These findings suggest that AimR and YosL act in concert with other yet unknown components to allow the switch from the lysogenic to the lytic cycle of SPβ.

The present study opens a wealth of interesting questions that need to be answered in the future. For instance, it remains to be resolved how the arbitrium system controls the activity of YopR. Moreover, it must be elucidated whether YopR serves as a transcription factor recognizing specific DNA elements or acts as a silencer to maintain the lysogenic state of SPβ. The latter possibility is not unlikely because the GC content of the SPβ genome is significantly lower than that of the *B. subtilis* genome [Lazarevic et al., 1999]. Furthermore, the function of YosL and other players of the lysis/lysogeny management system, such as YopN must be uncovered.

## SUPPLEMENTARY DATA

Supplementary Data are available Online.

## SOURCE DATA

Supplemental Dataset. The spreadsheet provides a summary of the conditions used for HDX-MS analyses and a full list of the peptides and residue-specific HDX obtained for YopR, YopR^G136E^ and YopR^Y304F^.

## DATA AVAILABILITY

Coordinates and structure factors have been deposited within the protein data bank (PDB) under the accession code: 8a0a. The authors declare that all other data supporting the findings of this manuscript are available within the article and its supplementary data files.

## ACKNOWLEDGEMENTS

We are grateful to Dr. Vladimir Lazarevic for providing the *B. subtilis* strain CU1147. We thank Pauline Plitzko for technical assistance. We also gratefully acknowledge the support of the Core Facility “Protein Spectroscopy” of the Medical School of the Philipps-University Marburg. We thank the European Synchrotron Radiation Facility (ESRF, Grenoble, France) and the Deutsche Elektronen Synchrotron (DESY, Hamburg, Germany) for excellent support.

## FUNDING

This work was supported by the DFG grants CO 1139/3-1 (to F.M.C.), BA 5311/9-1 in the framework of the SPP2330 “New Concepts in Prokaryotic Virus-host Interactions – From Single Cells to Microbial Communities” (jointly to G.B. and L.C), the core facility for interactions, dynamics and macromolecular assembly (project 324652314, to G.B.), and the Max Buchner Research Foundation (Re.3799 to R.H.).

## Conflict of interest statement

None declared.

---

**Figure S1.**
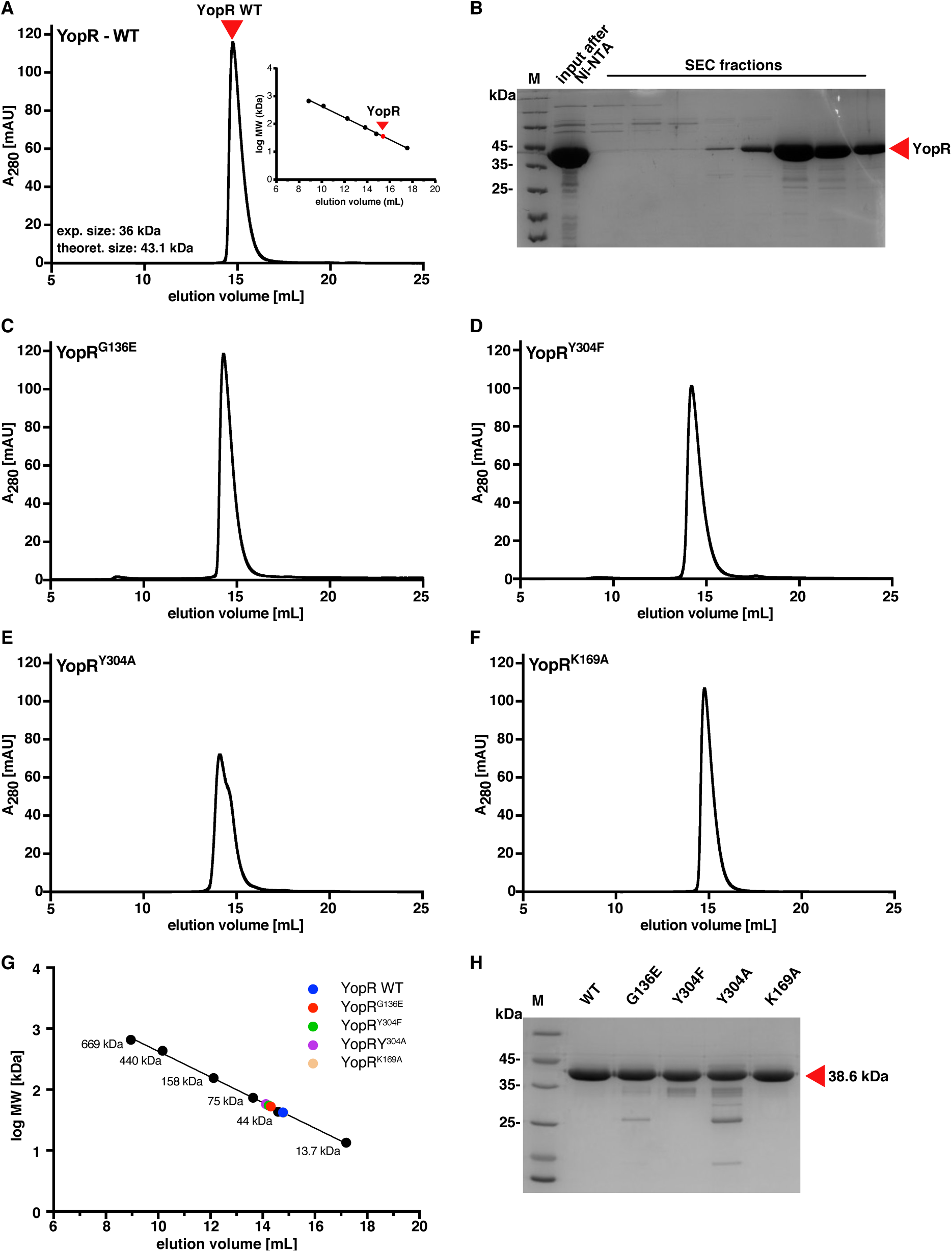
Analytical SEC and SDS PAGE analysis of YopR and YopR variants. **A.** Analytical SEC of YopR WT after additional heparin purification. Comparison to standard protein solutions is shown in the smaller graph. **B.** SDS-PAGE analysis of SEC fractions of YopR WT protein. **C-F.** Analytical SEC analysis of the variants YopR^G136E^ (**C**), YopR^Y304F^ (**D**), YopR^Y304A^ (**E**) and YopR^K169A^ (**F**). **G.** Comparison of YopR WT and variants to SEC of standard protein solutions (thyroglobulin [660 kDa], ferritin [474 kDa], aldolase [160 kDa], conalbumin [76 kDa], ovalbumin [43 kDa] and ribonuclease A [13.7 kDa]). **H.** SDS-PAGE of purified YopR WT, and the variants YopR^G136E^, YopR^Y304F^, YopR^Y304A^, and YopR^K169A^.

**Figure S2.**
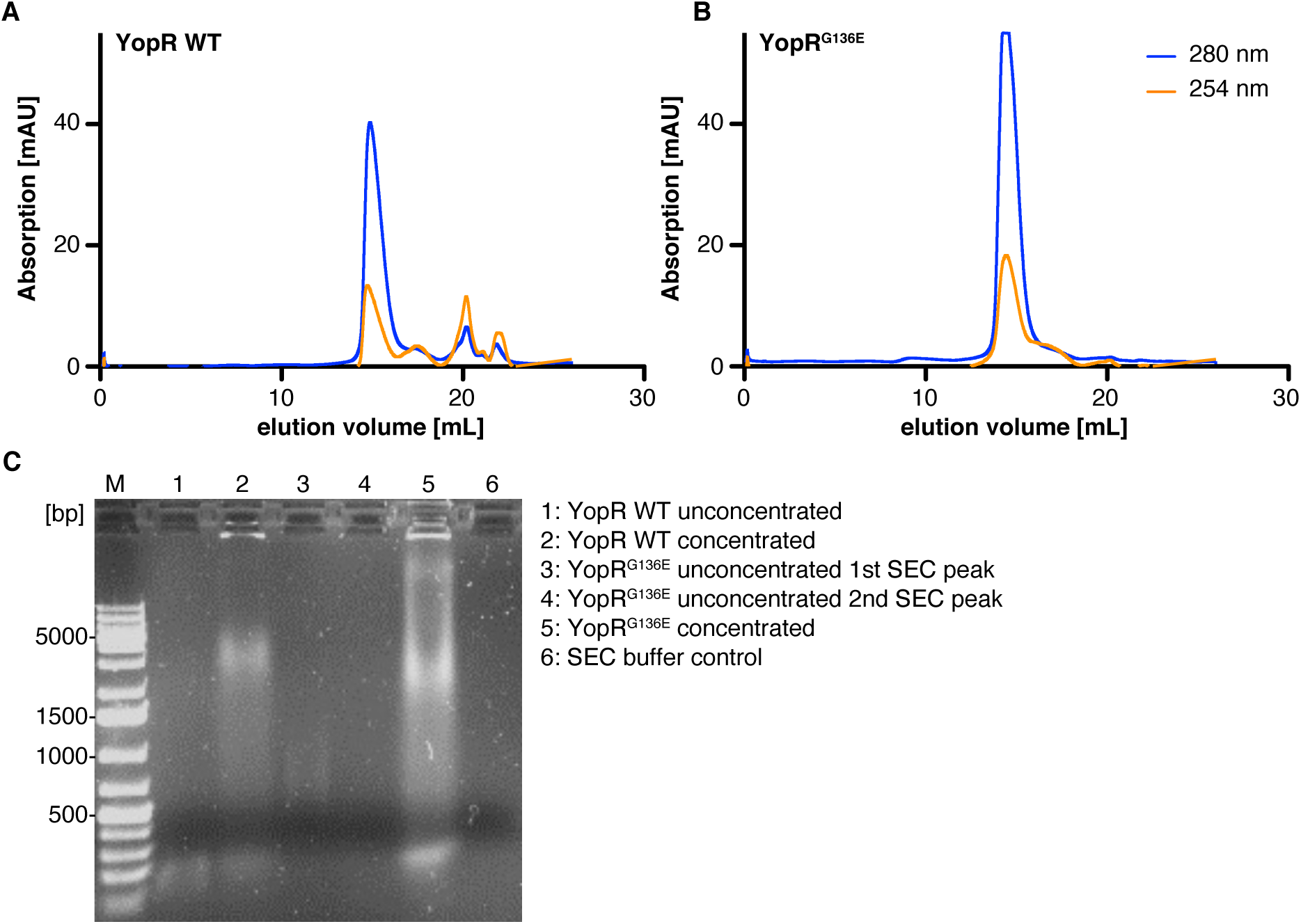
DNA contamination during purification of YopR and YopR^G136E^. **A, B.** Size-exclusion chromatography (SEC) of YopR WT (**A**) and YopR^G136E^ (**B**). Absorption at 280 nm is shown in blue, and 254 nm in orange. **C.** Analysis of co-purified DNA present in affinity purified YopR WT and YopR^G136E^ using agarose gel electrophoresis. Lanes are labelled within the figure.

**Figure S3.**
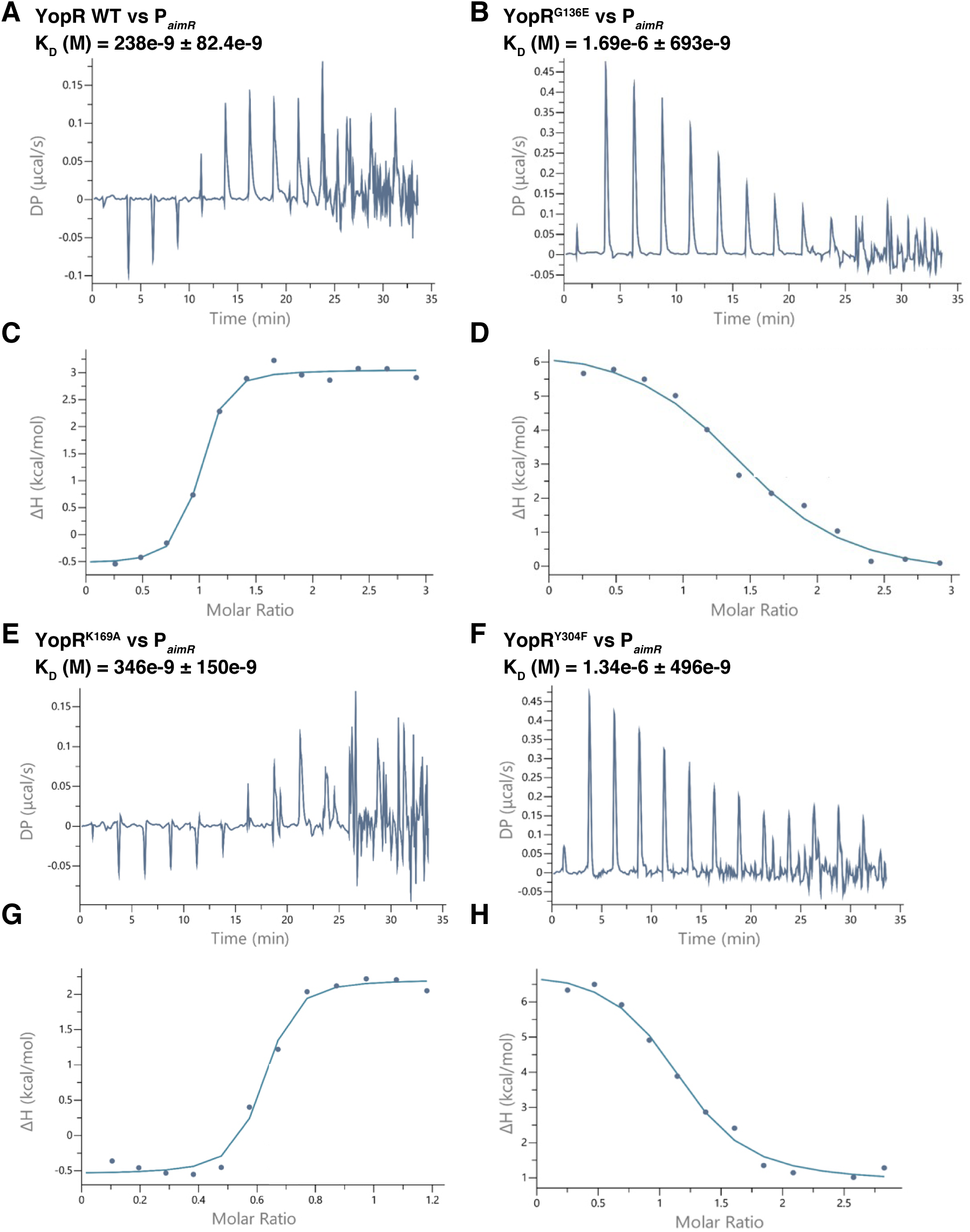
Binding of DNA to YopR and YopR variants determined by isothermal titration calorimetry (ITC). DNA of the *aimR* promoter region was added to the sample cell and titrated with purified YopR WT (**A**, **C**) and YopR variant proteins YopR^G136E^ (**B**, **D**), YopR^K169A^ (**E**, **G**) and YopR^Y304F^ (**F**, **H**). Panels A, B, E and F show the thermograms for each ITC run, and panels C, D, G and H are the resulting fitted plots; the blue dots represent the binding enthalpies (ΔH) per injection versus ligand concentration. DP stands for differential power.

**Figure S4.**
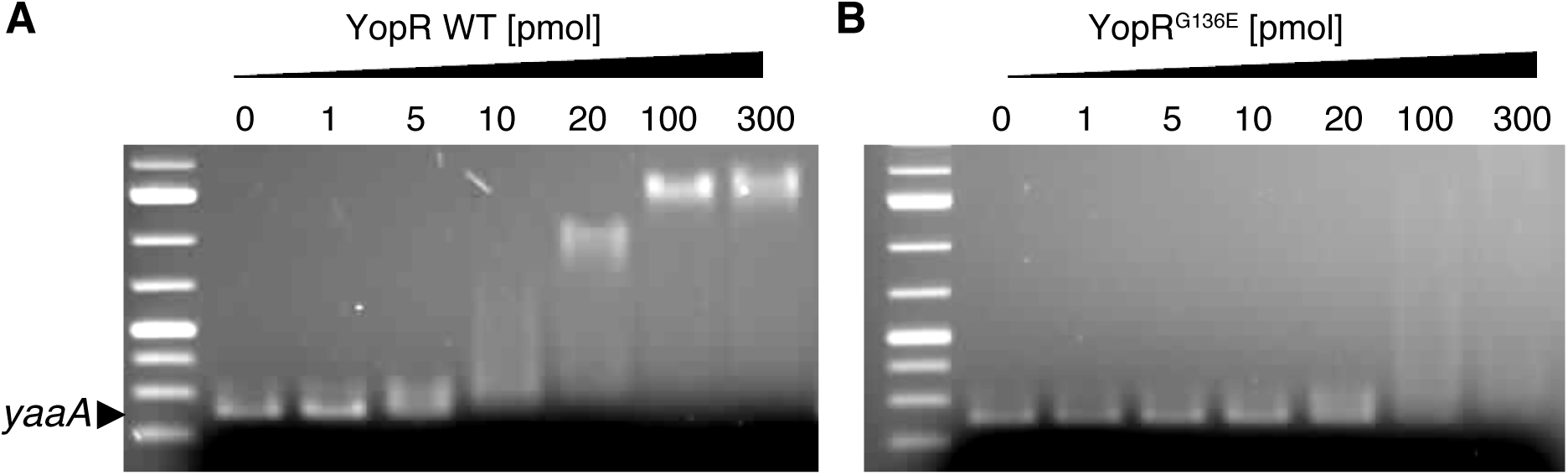
Electrophoretic mobility shift assays (EMSAs) showing the DNA-binding activities of the YopR wild type protein and the YopR^G136E^ variant. **A, B**. Complex formation between the YopR wild type protein (**A**) and the YopR^G136E^ variant (**B**) with the unrelated *B. subtilis* DNA fragment *yaaA*.

**Figure S5.**
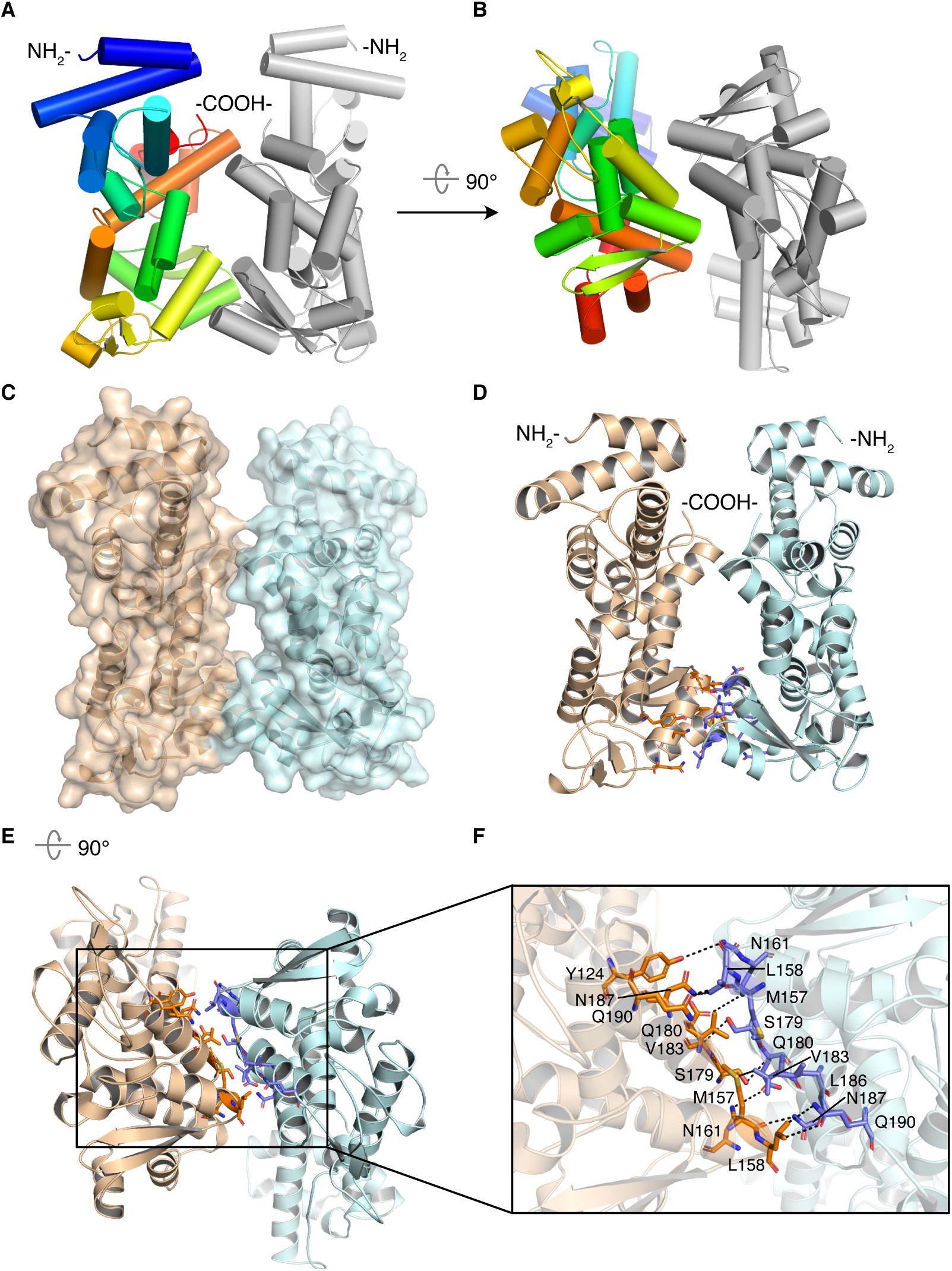
Analysis of the asymmetric unit (ASU) of the YopR crystal structure. **A, B.** Overall fold of the two YopR molecules found in the asymmetric unit (ASU) of the crystallographic analysis. One monomer is shown in rainbow color and one in grey. **C.** Surface representation of the two monomers present in the ASU shown in beige and light blue, respectively. **D.** Interacting residues in helices H’ and I of the two YopR monomers are shown as orange and purple sticks. **E, F.** Zoom into the region of helices H’ and I highlighting the interacting residues found in the ASU of the YopR crystal structure.

**Figure S6.**
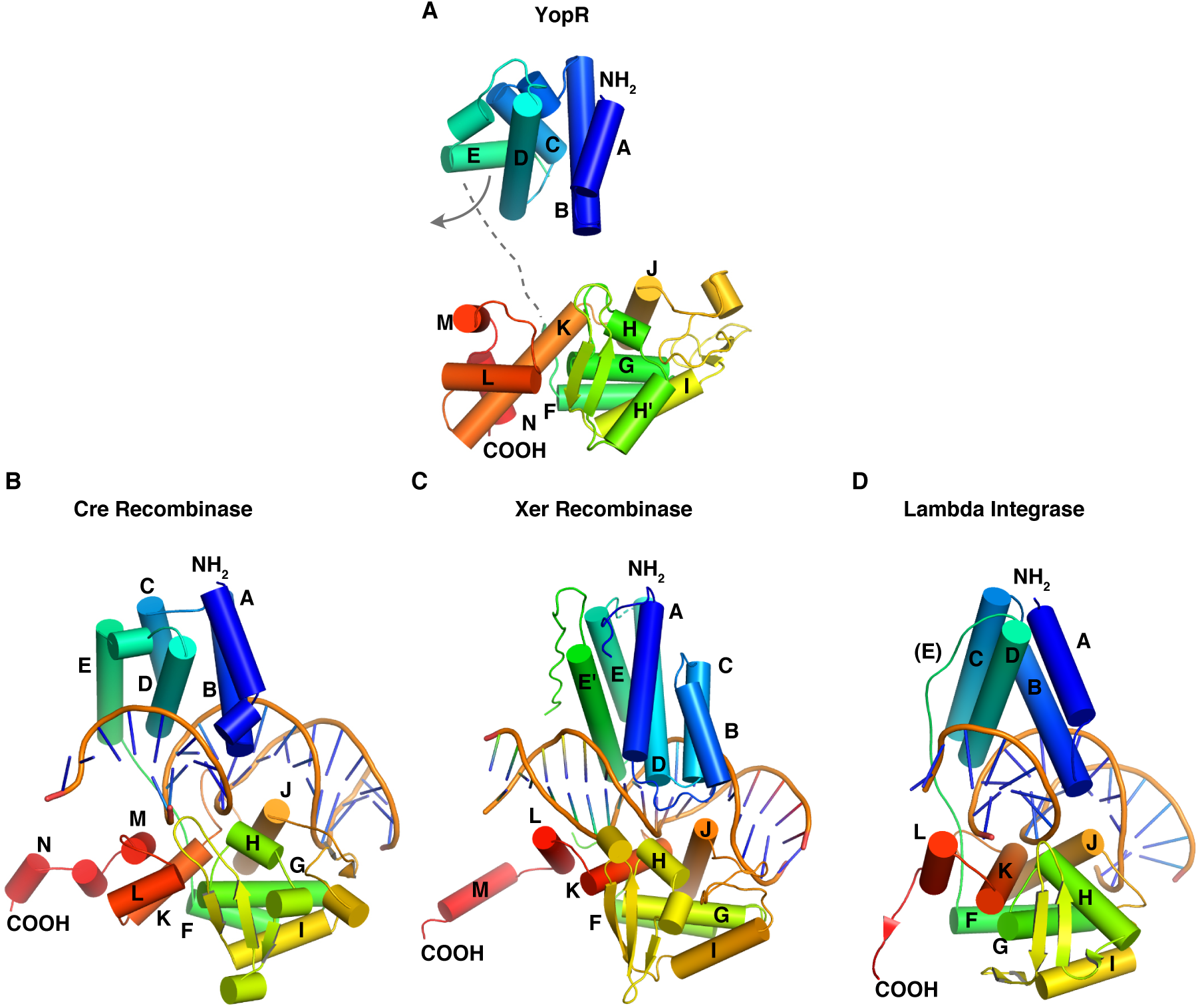
Structural comparison of YopR to known tyrosine recombinases. **A.** N- and C-terminal domains of the YopR crystal structure were individually overlayed with the structure of the Cre recombinase Cre (PDB-ID: 1q3u) [Ennifar et al., 2003] as described in the main text and figure 5. **B-D.** Crystal structures of the tyrosine recombinases (**B**) Cre (PDB-ID: 1q3u) [Ennifar et al., 2003]; (**C**) XerH (PDB-ID: 5jk0) [Bebel et al., 2016], and (**D**) Lambda Int (PDB-ID: 1p7d) [Aihara et al., 2003], all bound to DNA.

**Figure S7.**
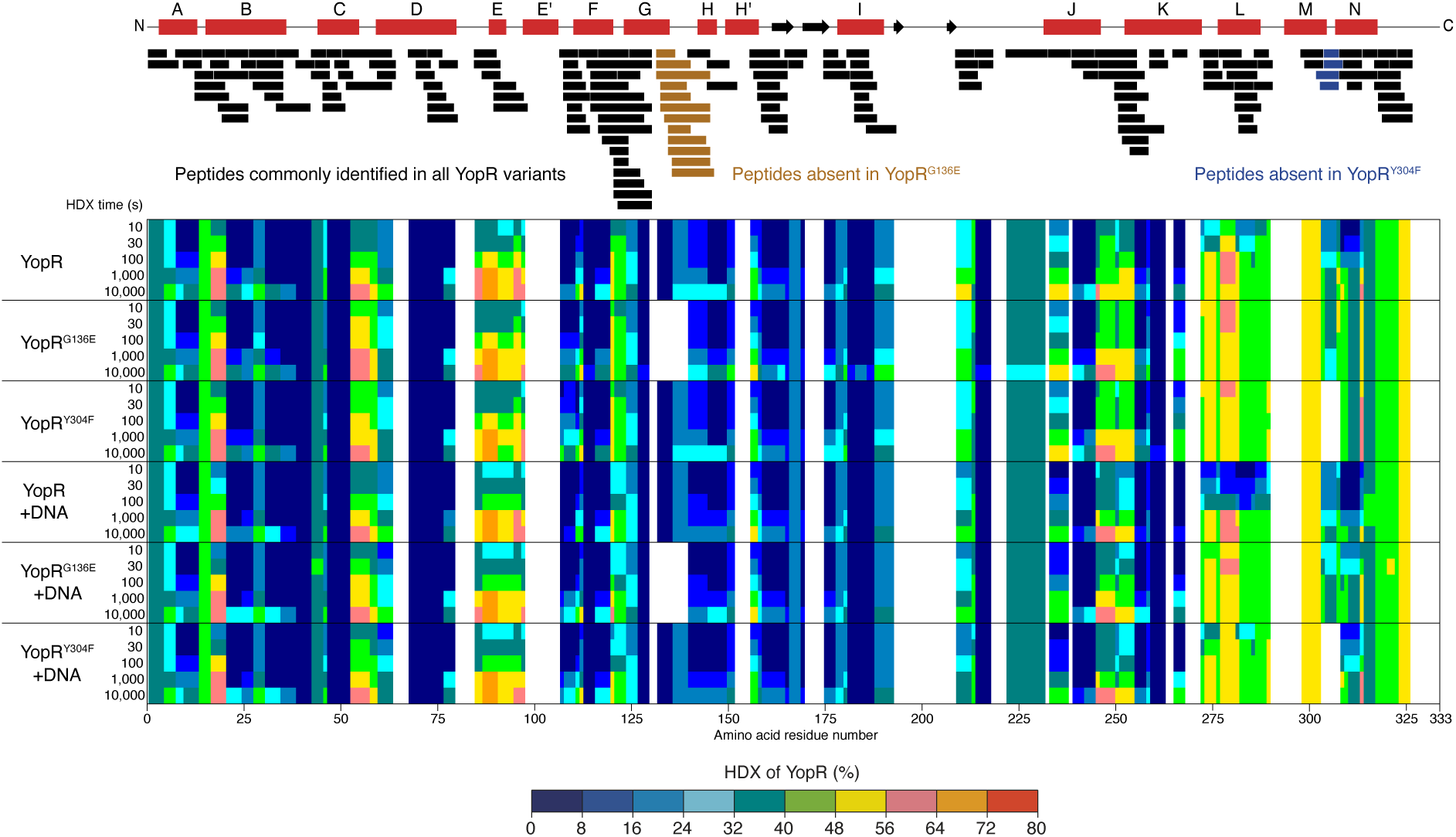
Hydrogen/deuterium exchange mass spectrometry of YopR. Each bar represents a peptide of YopR that was identified in HDX-MS experiments. Peptides that could not be identified for YopR^G136E^ or YopR^Y304F^ variants due to the variations are highlighted in ocre and blue, respectively. The secondary structure of YopR is depicted above (red boxes, *α*-helices; black arrows, β-strands). The residue-specific HDX of individual YopR, YopR^G136E^ and YopR^Y304F^ residues in apo and DNA-bound states was calculated with DynamX 3.0 from peptides as follows: When any residue was covered by only a single peptide, the residue-specific deuterium uptake equals that of the whole peptide. In the case of overlapping peptides for any given residue, residue-specific deuterium uptake was determined by the shortest peptide covering that residue. Where multiple peptides were of the shortest length, the peptide with the residue closest to the peptide C-terminus was utilized. Numerical values for deuterium uptake by peptides and residue-specific HDX are provided in Supplemental Dataset.

**Figure S8.**
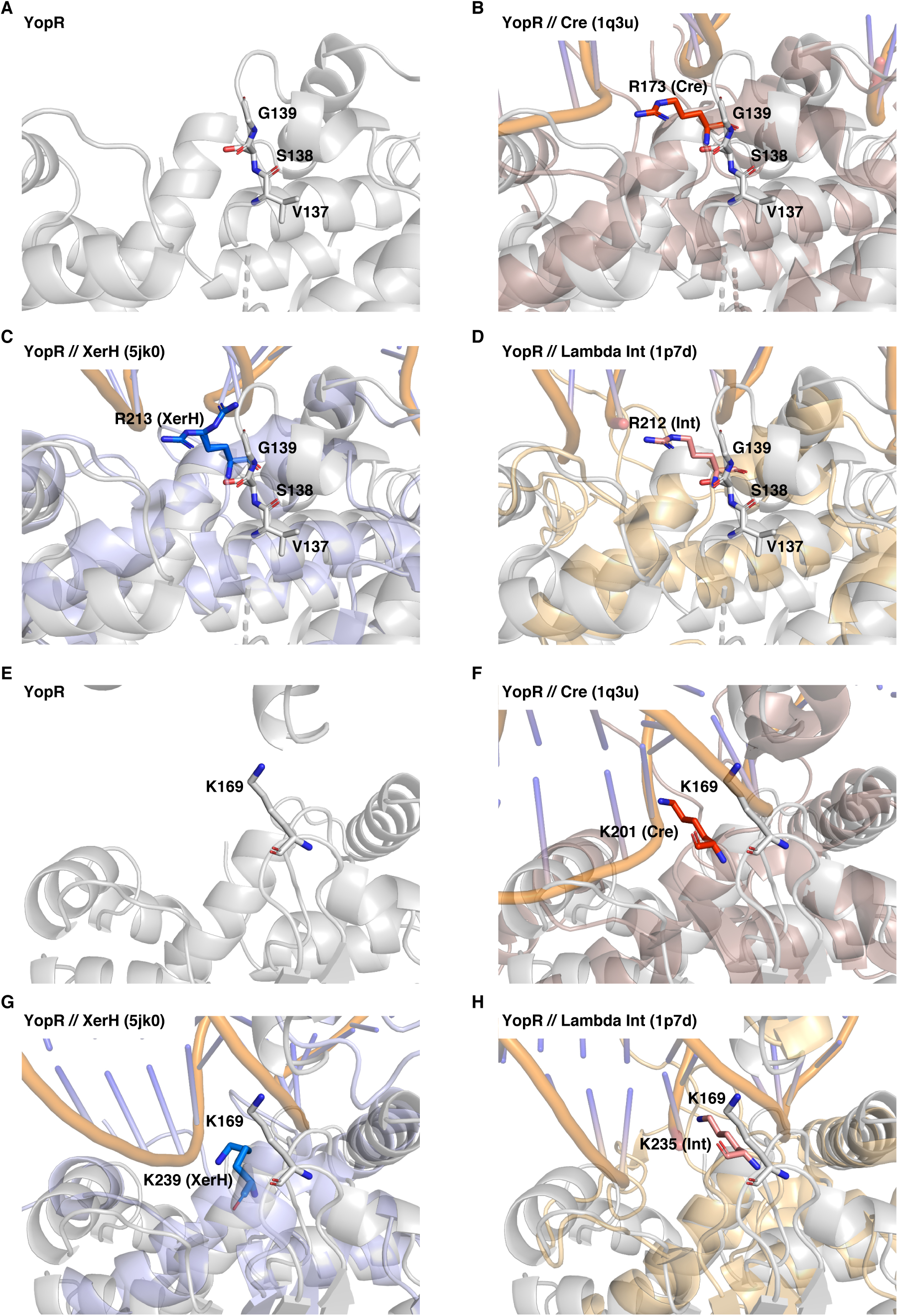
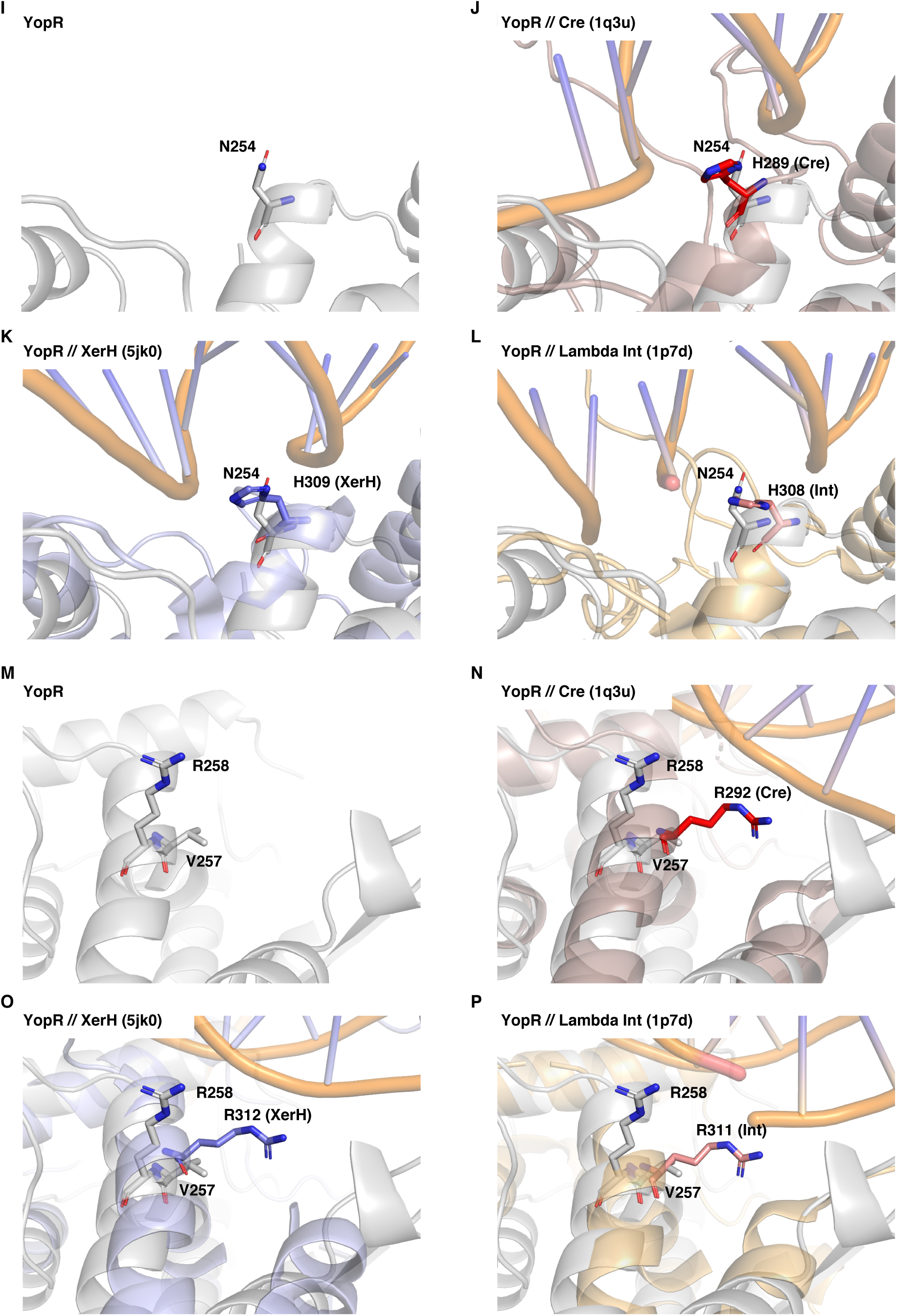
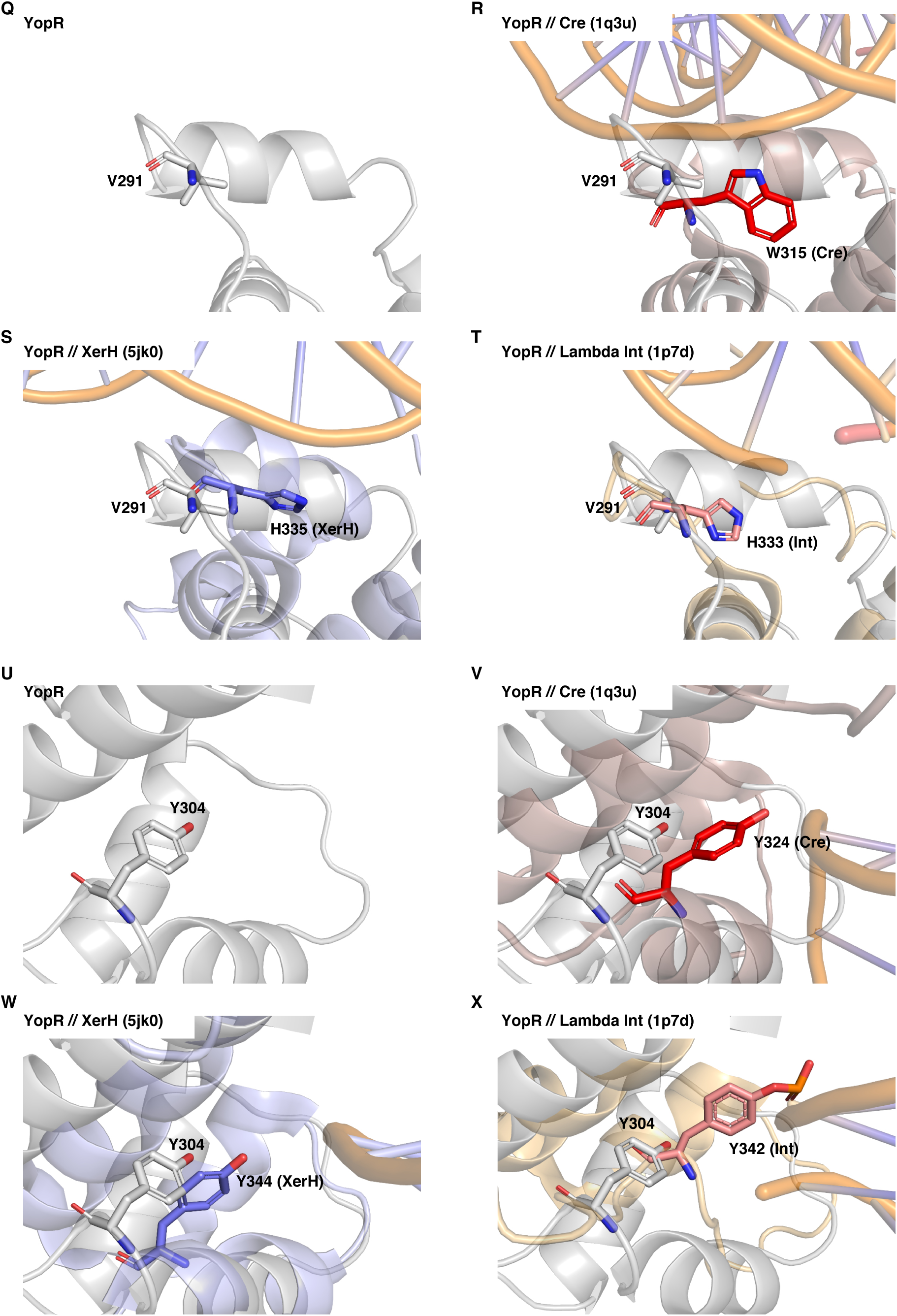
Structural comparison of catalytically important amino acid residues of tyrosine recombinases to those found in YopR. **A-D.** Comparison of V137/S138/G139 in YopR (**A**) to R173 in Cre (**B**), R213 in XerH (**C**) and R212 in Lambda Int (**D**). E-H. Comparison of K169 in YopR (**E**) to K201 in Cre (**F**), K239 in XerH (**G**) and K235 in Lambda Int (**H**). I-L. Comparison of N254 in YopR (**I**) to H289 in Cre (**J**), H309 in XerH (**K**) and H308 in Lambda Int (**L**). M-P. Comparison of V257/R258 in YopR (**M**) to R292 in Cre (**N**), R312 in XerH (**O**) and R311 in Lambda Int (**P**). Q-T. Comparison of V291 in YopR (**Q**) to W315 in Cre (**R**), H335 in XerH (S) and H333 in Lambda Int (T). U-X. Comparison of Y304 in YopR (U) to Y324 in Cre (V), Y344 in XerH (W) and Y342 in Lambda Int (X).

**Figure S9.**
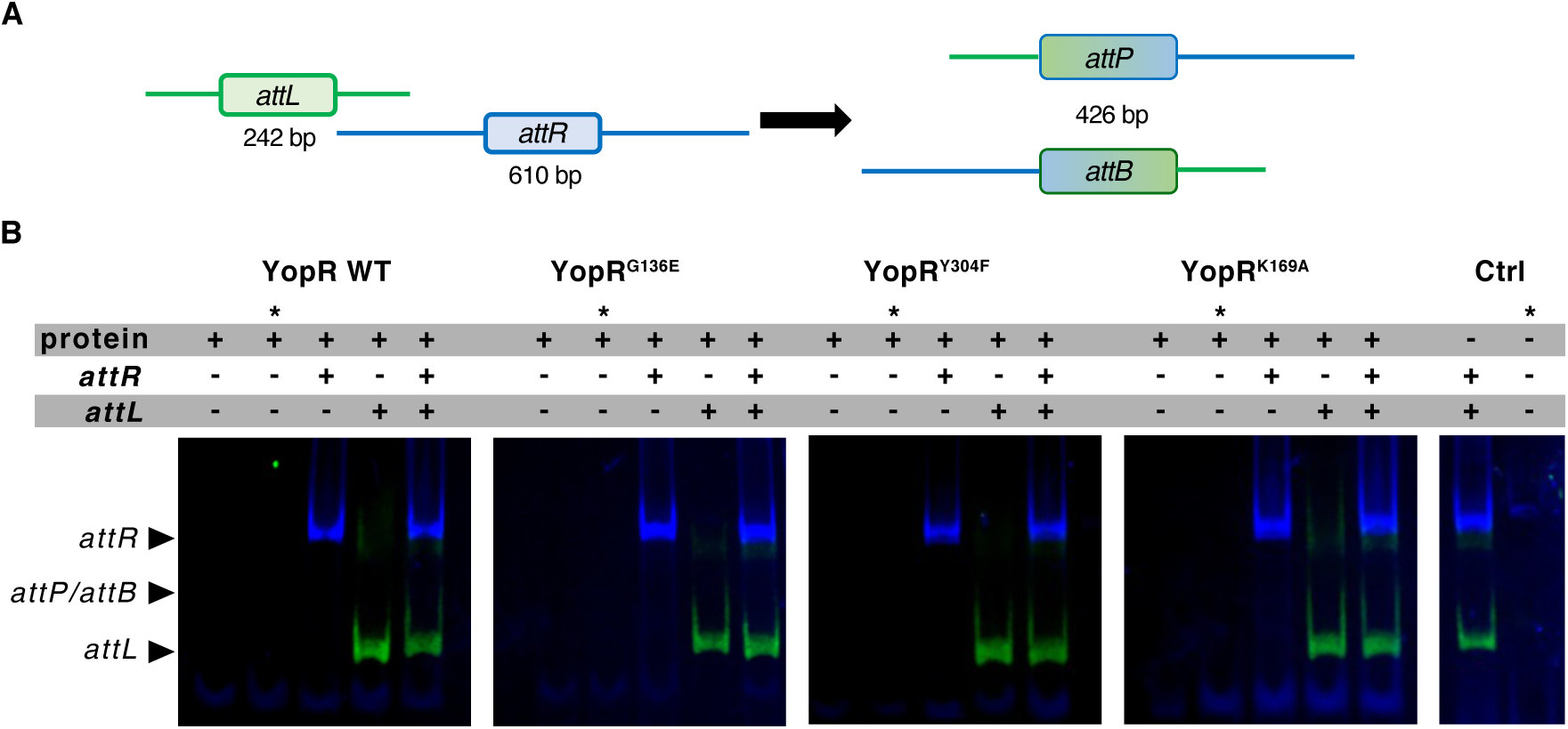
*In vitro* recombinase assay. **A.** Schematic overview of the *in vitro* recombination. The *attL* fragment (242 bp) is labeled with Cy5 (green) and the *attR* fragment (610 bp) is labeled with Cy3 (blue). The resulting hybrid *attP* and *attB* fragments indicative for DNA recombination have an expected size of 426 bp. **B.** Recombinase assay using the *attR* and *attL* regions of the SPβ phage [Abe et al., 2017] assessing the recombinase activity of the purified YopR WT protein and the YopR variants YopR^G136E^, YopR^Y304F^, and YopR^K169A^. Lanes marked with an asterisk (*) are reactions in which the labeled DNA fragments were exchanged by an unlabeled DNA fragment carrying no fluorescent dye.

**Table S1.**
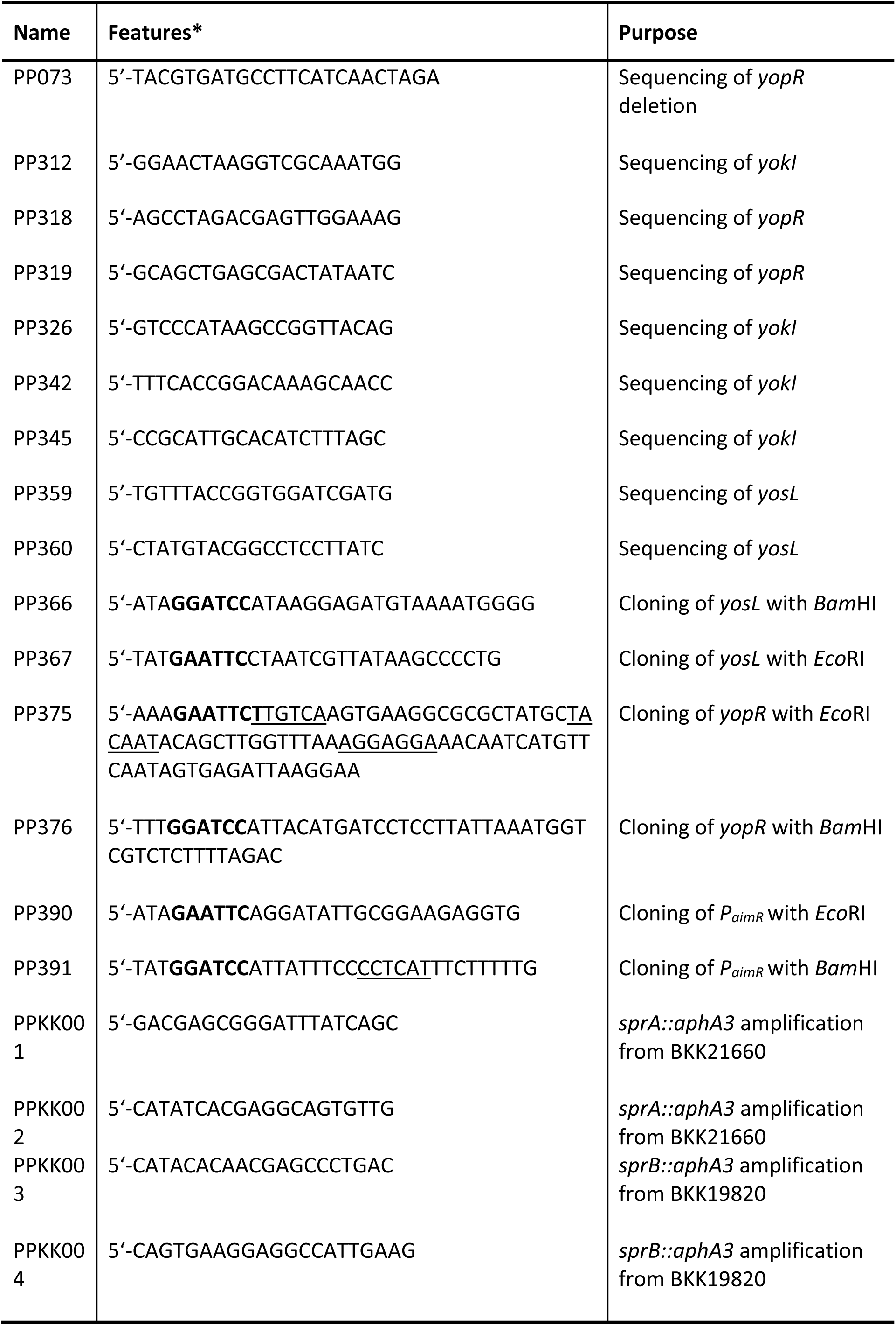
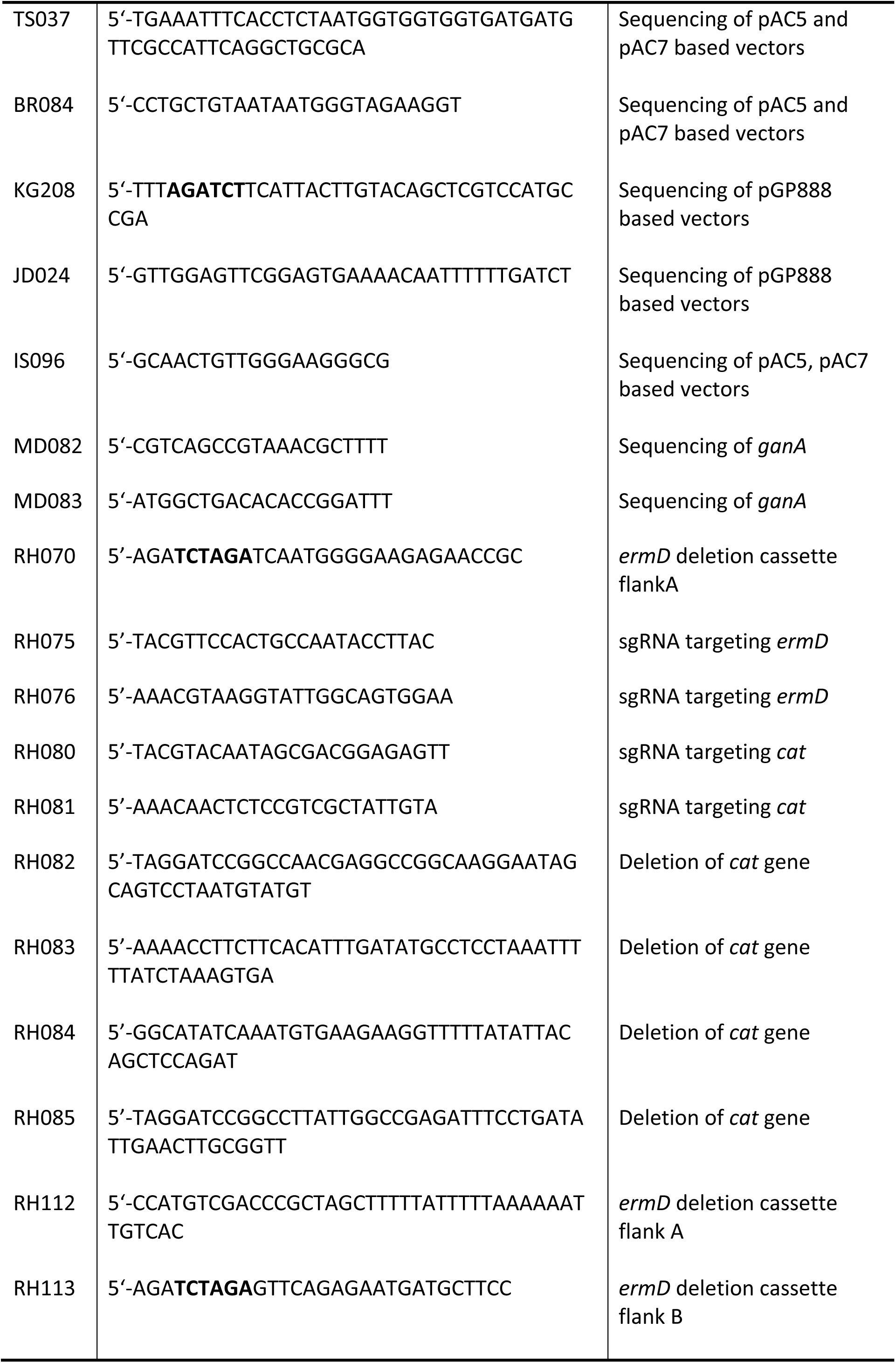
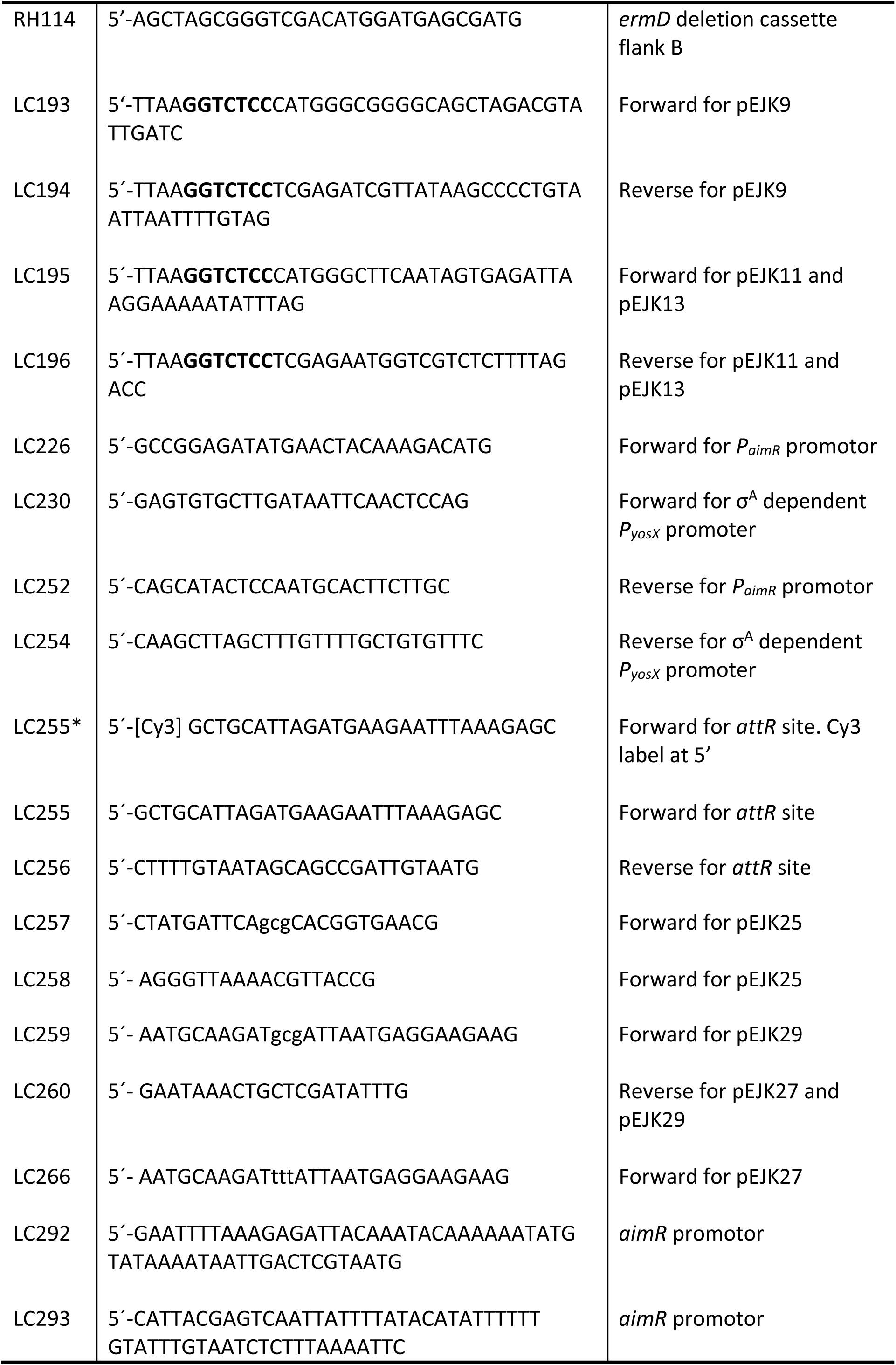
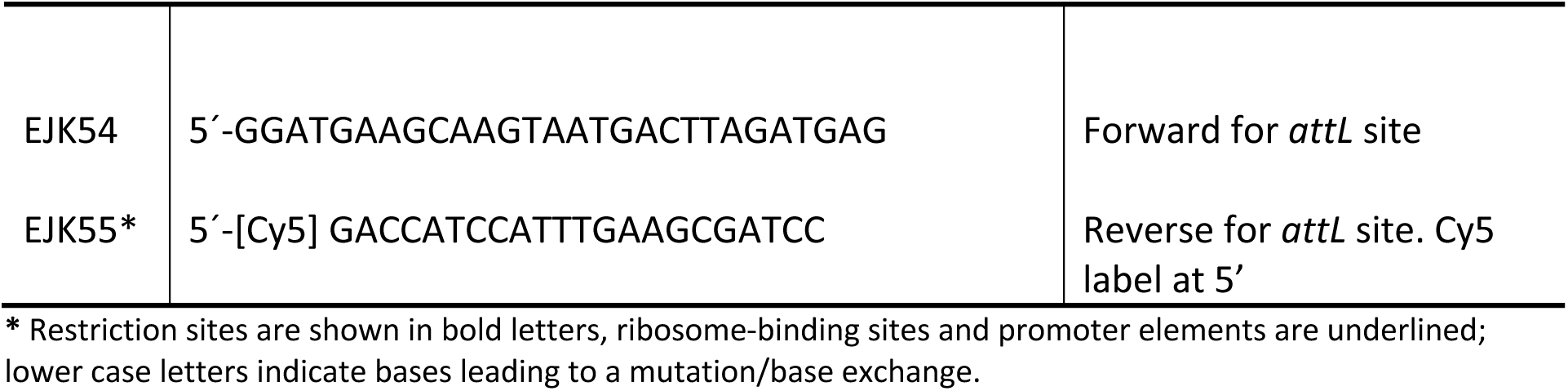
Primers used in the study.

**Table S2.**
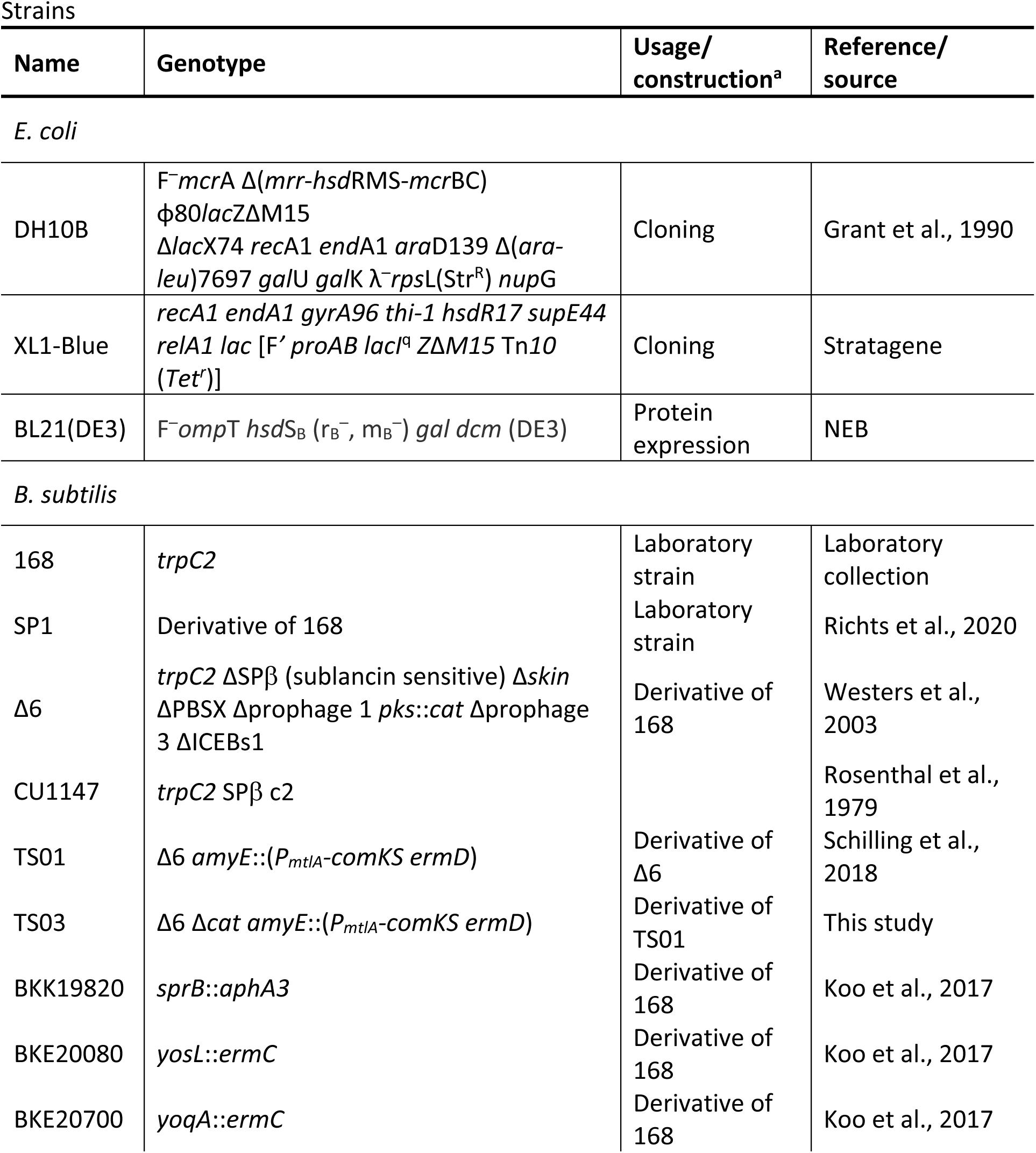
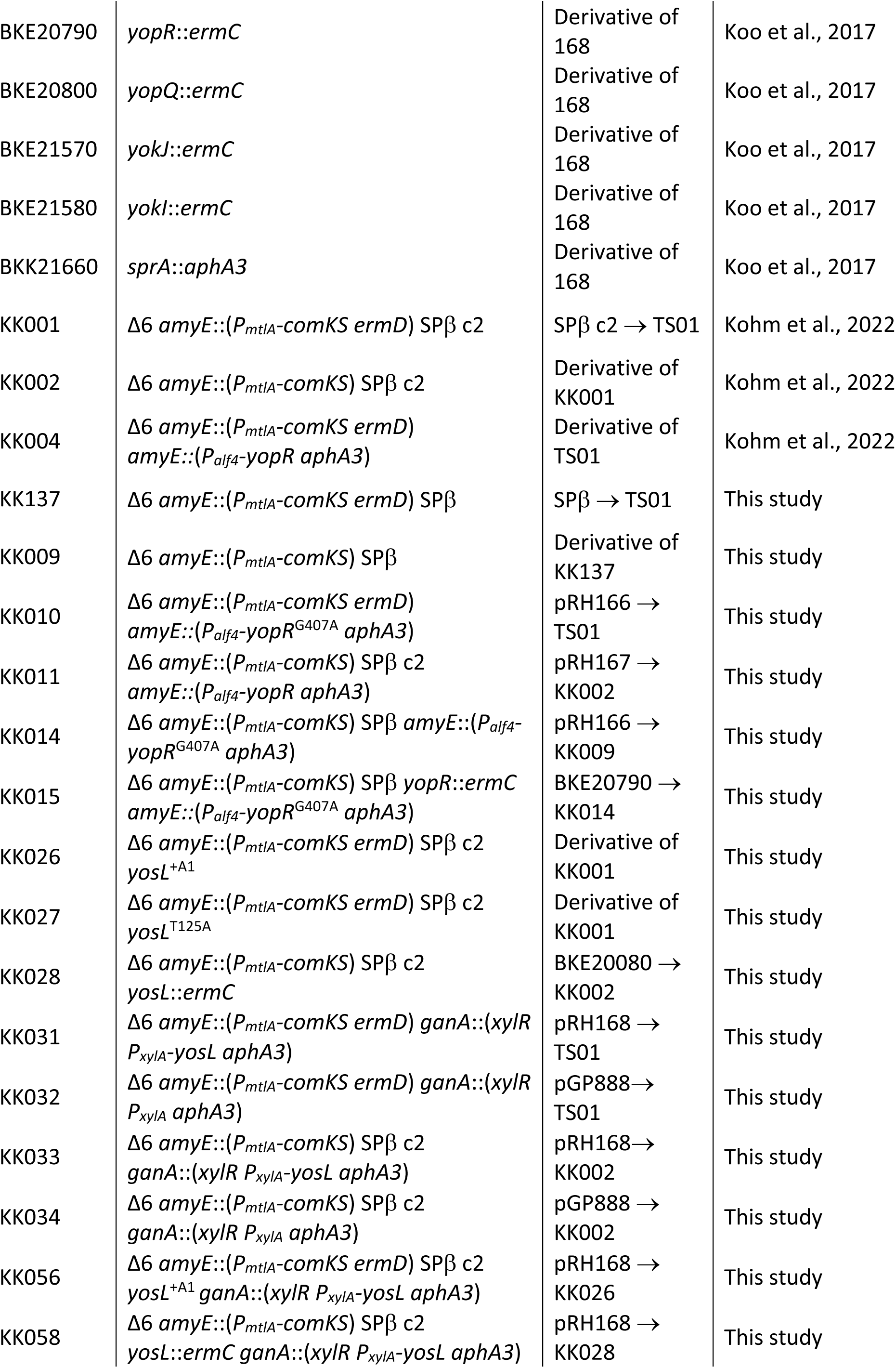
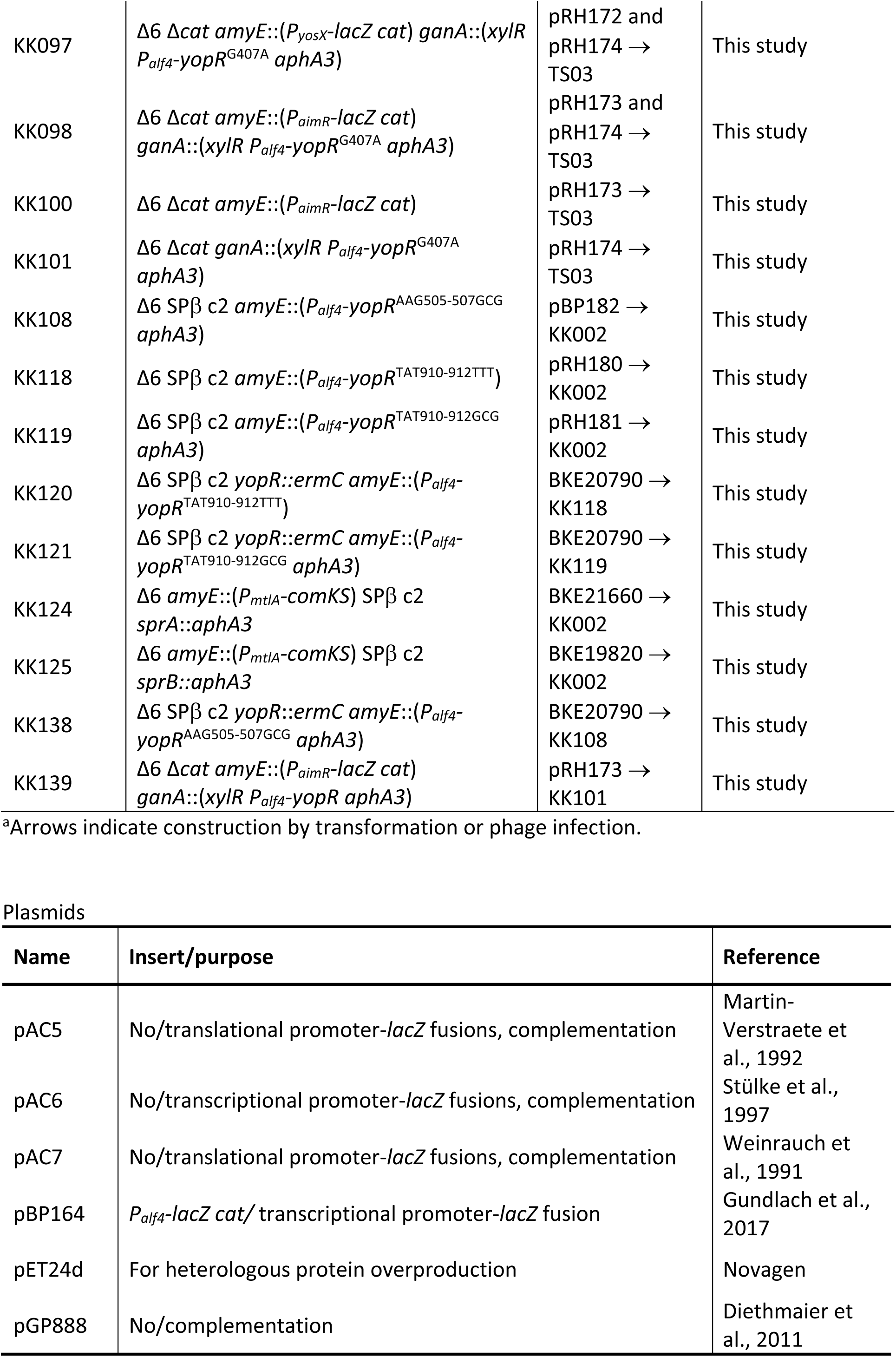
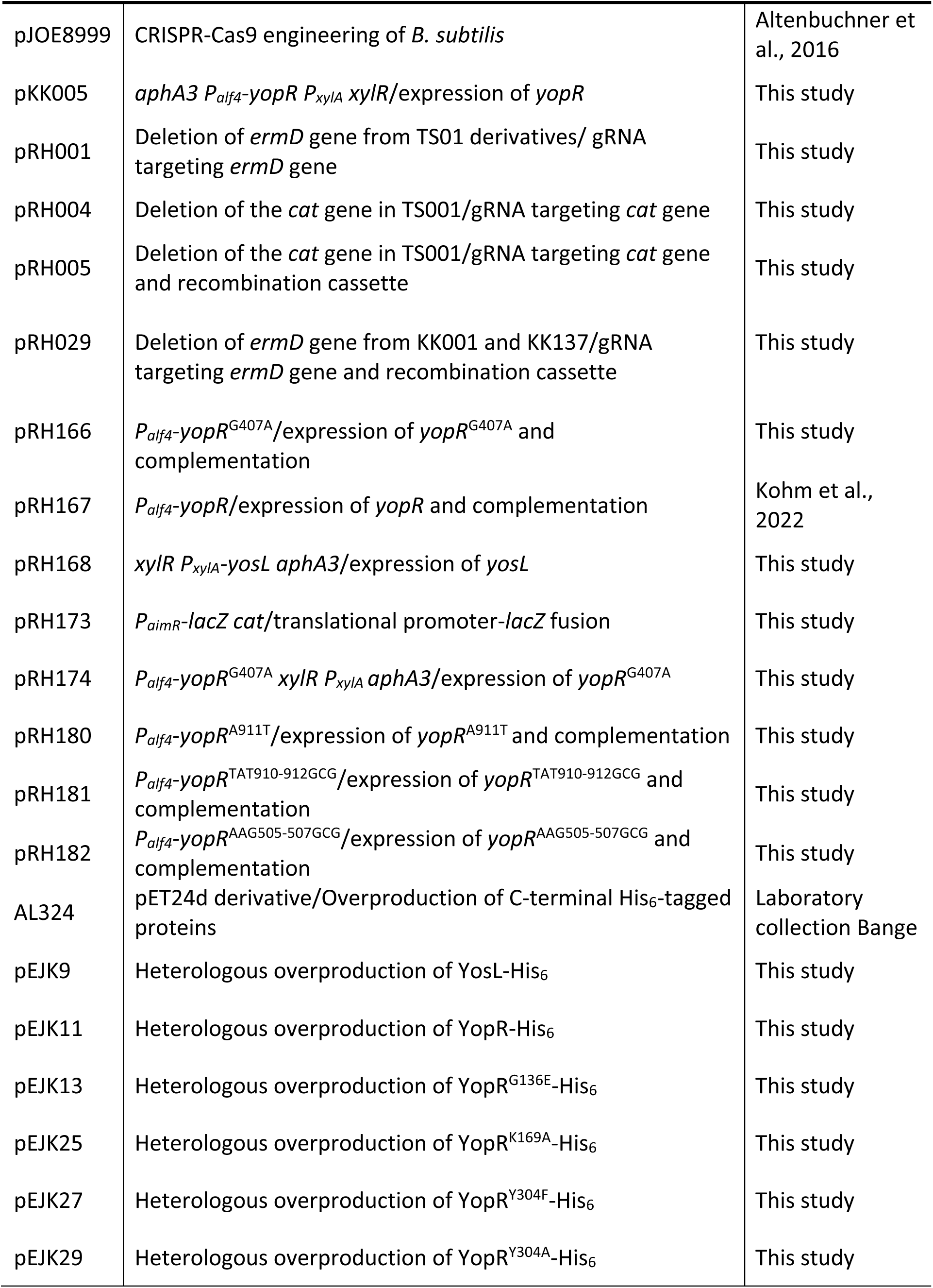
Strains and plasmids

## Notes

### Competing Interest Statement

The authors have declared no competing interest.

### Summary of Updates

The previous version of the manuscript contained few typos. Moreover, we performed additional control experiments and repeated a promoter activity assay. Furthermore, HDX experiments were expanded to assess the DNA-binding activity of the SPbeta repressor YopR.

